# Seasonal and interannual variation of common dolphin’s densities in Portuguese waters

**DOI:** 10.1101/2025.04.16.649110

**Authors:** Miguel P. Martins, Marc Fernandez, Ana Marçalo, Nuno Oliveira, Tiago A. Marques

## Abstract

Modelling a species ecology and abundance provides important insights into its habitat preferences, population trends and distribution. This can be used to inform conservation efforts, highlighting populations at risk of extinction and regions in need of protection. Here, we studied how environmental factors relate to common dolphin’s (*Delphinus delphis*) densities in the waters off mainland Portugal. We analyzed an opportunistic dataset collected using European Seabirds At Sea (ESAS) methodology with distance sampling, spanning from 2004 to 2020. There were 1151 sightings of common dolphins, 737 of which were during effort. Detection functions were fitted to model the dolphin’s detectability as a function of the distance to the track line. We used smooths of static (depth, slope, and distance to the coast, 200m and 1000m isobaths) and dynamic (sea surface temperature, salinity, chlorophyll-a concentration with and without a time lag, zooplankton, and sardine biomass) environmental factors, relating them to counts corrected for detectability and survey effort, to fit habitat-based Density Surface Models. We then predicted and mapped common dolphin densities seasonally (winter, spring, summer autumn) throughout the sample years. The results show how different environment variables relate to dynamic common dolphin densities. In general, common dolphins had higher densities in coastal environments, especially during the summer due to high productivity and lower sea surface temperature in shelf waters, associated with seasonal upwelling. During the summer, there were predicted local peaks in common dolphin densities, in areas with high zooplankton and chlorophyll-a concentration. In the other seasons, its range expands to more offshore waters, with less local density peaks. The interannual variation of this species’ densities is mostly explained by the annual sardine biomass. Overall, this brings important insights into this species conservation and potential impacts from anthropogenic activities, such as climate change and overfishing.

## 1 INTRODUCTION

Estimating abundance, trends, and species distributions, can contribute to population conservation and management. Despite this, there are still great gaps in our knowledge of cetacean densities, with large areas still not surveyed or with not enough effort to allow for trend analysis (Kaschner et al., 2012). Distance sampling is a widely used data collection methodology to achieve unbiased abundance and density estimates. By recording the distance an object is from the trackline, one can fit a model that gives the detection probability conditional to the distance, i.e. a detection function, allowing to correct the number of animals sighted for their detectability (Buckland et al., 2001). The detection function can be further improved by accounting for other covariates besides distance that also impact detectability (Marques et al., 2007). Several distance sampling methods can be employed, like point counts, quadrat sampling, strip transect sampling and line transect sampling (Buckland et al., 2001). Line transect sampling has been repeatedly used for cetacean abundance studies (Kaschner et al., 2012). Transects can be split into segments and counts per segment can then be modelled to produce abundance estimates conditional to environmental variables (Miller et al., 2013). The model results can then be projected in space, creating spatially explicit density estimates that reflect the animal’s distribution (Miller et al., 2013; Roberts et al., 2016). These models are referred to as Density Surface Models (DSMs), which can be approached either by a one-stage (when the parameters of the detection function and the spatial/environmental model are estimated at the same time) or a two-stage approach (when the detection function and the spatial/environmental model are fitted separately) (Miller et al., 2013).

As of 2012, about 25% of the global ocean surface had been covered by line transect surveys dedicated to studying cetacean abundance (Kaschner et al., 2012). Since then, there has been greater sampling effort worldwide, for example: (Bedriñana-Romano et al., 2022; Carretta et al., 2024; Gilles et al., 2023; Pike et al., 2019), however there are still large areas of the ocean that haven’t been properly surveyed. The NASS and SCANS surveys have brought insight into the summer abundance and distribution of cetacean species in the eastern and central north Atlantic (Gilles et al., 2023; Pike et al., 2019). Despite large, dedicated surveys not occurring yearly, sampling effort in European Atlantic waters has been extensive enough to allow for some trend analysis, although, effort has been spatially uneven throughout the years. There has been notably less effort on the west and south waters of mainland Portugal, where sampling effort has been restricted to coastal areas until only the last SCANS survey, where it was expanded offshore (Gilles et al., 2023; Hammond et al., 2021). Additionally, sampling effort is typically restricted to the summer, in detriment to the other seasons. Therefore, we have an incomplete understanding of the seasonal and interannual variation of cetaceans’ abundance and distribution in the region, especially in offshore waters. Portuguese waters are included in the Canary/Iberian Eastern Boundary Upwelling System (Bakun et al., 2015), among the most productive marine ecosystems in the world. This makes the area suitable for the occurrence of marine megafauna (Couto et al., 2017, 2018; Martins et al., 2025). This region is of particular interest to study cetaceans, having 27 confirmed species recorded, through sightings and strandings (Mathias et al., 2024; Morais et al., 2021).

Despite lower survey effort, when compared with other European waters, it is clear that the common dolphin (*Delphinus delphis*) is the most abundant cetacean in Portugal, with the most recent abundance estimate at 76,615 individuals over an area of 210,991 km² (Gilles et al., 2023). This results in an average density of 0.363 individuals/km² in Portuguese waters. However, this estimate is for only about 68% of the total Portuguese Exclusive Economic Zone (EEZ) - here defined as the sum of the area of the EEZ itself, territorial sea and inland maritime waters (*DGRM*, 2018). This globally distributed species has a patchy distribution along Portuguese coastal waters, with its occurrence being largely related to high primary production (Moura et al., 2012). In fact, its occurrence has been associated with upwelling modified habitats (Au & Perrymanl, 1985), which create suitable conditions for the small pelagic fishes the dolphin targets, like the sardine (*Sardina pilchardus*) – the most important prey item on the common dolphin’s diet in Portugal (Marçalo et al., 2018). The common dolphin tends to prefer coastal areas (Gilles et al., 2023), although its presence is also detected in offshore environments in lower densities (Gilles et al., 2023). Despite occurring in both tropical and temperate waters, the common dolphin’s habitat suitability has been related to relatively low (<20° C) sea surface temperature (SST), in the Eastern North Atlantic (Fernandez et al., 2021).With climate change, it is expected that the common dolphin’s and its preferred prey’s range expands toward higher latitudes (Lima et al., 2022; MacLeod, 2009).

In this study, we model common dolphin densities from an opportunistic line transect dataset, through habitat-based DSMs. We also explore potential climate change and overfishing impacts on the common dolphin’s densities.

## 2 METHODS

### 2.1 Data collection

From December 2004 to December 2020, line transects with two observers were employed by SPEA (Sociedade Portuguesa para o Estudo das Aves) for marine bird data collection. In total, 62,392.6 km were surveyed. Survey effort covered extensively mainland Portugal’s shelf waters, with 75% of the searched transects having taken place in waters shallower than 220 m deep. Still, offshore areas were surveyed, however, with much lower intensity (Figure 1). Distance sampling adapted from (Tasker et al., 1984) was opportunistically employed whenever cetaceans were sighted. Cetaceans were recorded on one side of the boat and sightings were considered on effort when the distance to the transect line was inferior to 300 m. Distance to the line was binned, during data collection, into five categories: i) 0-50 m; ii) 50-100 m; iii) 100-200 m, iv) 200-300 m, v) >300 m. Sightings with a distance at detection greater than 300 m were recorded but considered as outside sampling effort and, therefore, removed from the analysis.

**FIGURE 1.**
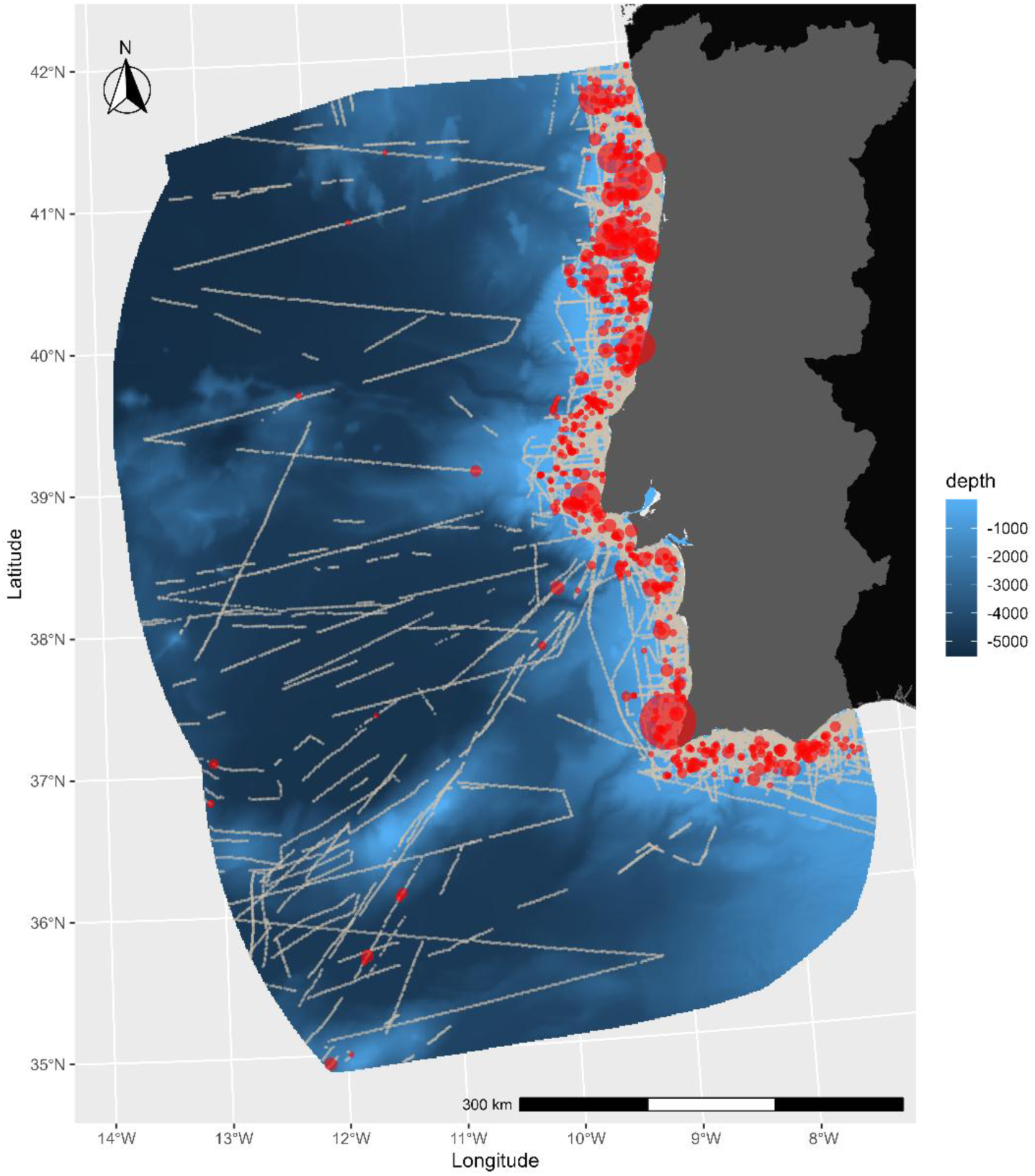
Transect effort (light grey) and common dolphin sightings on effort (red) recorded from 2004 to 2020 in mainland Portugal’s EEZ. The common dolphin sightings point size is proportional to the group size.

### 2.2 Environmental variables

A set of 11 environmental variables was used for model fitting and predictions (Table 1). Five static variables were considered (seafloor depth, seafloor slope and distances to the coast, to the 200 m and 1000 m isobathymetric lines). These variables were derived from an esriAscii file containing a digital terrain model (DTM) for bathymetry (seafloor depth at a given location), downloaded from the GEBCO initiative on 30/09/2024 (GEBCO, 2025). Depth was read directly from the DTM in R, version 4.4.1 (R Core Team, 2024) using the raster package (Hijmans, 2023). From the DTM file, the seafloor slope was calculated in R using the terrain function from the raster package (Hijmans, 2023). Distances to the coast and to the 200 m and 1000 m isobathymetric lines were derived from the DTM using the dist2isobath function from the marmap R package (Pante et al., 2023)

**TABLE 1.** Variables used for modelling and predicting common dolphin densities. NA (non-applicable); 8D (8-day composite); Y (yearly).

| Variable | Acronym/<br>Abbreviation | Temporal<br>resolution | Spatial<br>resolution | Units | Source/Product ID |
| --- | --- | --- | --- | --- | --- |
| Seafloor depth | Depth | NA | 15 arcsec | m | GEBCO |
| Seafloor slope | Slope | NA | 15 arcsec | ° | GEBCO |
| Distance to the coast | DTC | NA | NA | m | GEBCO |
| Distance to the 200 m isobathymetric line | DT200 | NA | NA | m | GEBCO |
| Distance to the 1000 m isobathymetric line | DT1000 | NA | NA | m | GEBCO |
| Sea surface temperature | SST | 8D | 0.083° | °C | Atlantic-Iberian Biscay Irish- Ocean Physics Reanalysis.<br>Product identifier: IBI_MULTIYEAR_PHY_005_002 |
| Chlorophyll-a concentration | CHL | 8D | 0.083° | mg/m<sup>3</sup> | Atlantic-Iberian Biscay Irish- Ocean BioGeoChemistry NON ASSIMILATIVE Hindcast.<br>Product identifier: IBI_MULTIYEAR_BGC_005_003 |
| Chlorophyll-a concentration with one-month lag | CHL-lag | 8D | 0.083° | mg/m<sup>3</sup> | Atlantic-Iberian Biscay Irish- Ocean BioGeoChemistry NON ASSIMILATIVE Hindcast.<br>Product identifier: IBI_MULTIYEAR_BGC_005_003 |
| Salinity | Salinity | 8D | 0.083° | 10<sup>-3</sup> | Atlantic-Iberian Biscay Irish- Ocean Physics Reanalysis.<br>Product identifier: IBI_MULTIYEAR_PHY_005_002 |
| Mass content of zooplankton expressed as carbon in sea water | Zooplankton | 8D | 0.083° | g/m<sup>2</sup> | Global ocean low and mid trophic levels biomass content hindcast.<br>Product identifier: GLOBAL_MULTIYEAR_BGC_001_033 |
| Iberian shelf waters Spring sardine biomass | Biomass | Y | NA | tonnes | ICES 2024 |

Additionally, six dynamic variables were considered (chlorophyll-a concentration – with and without a one-month time lag, sea surface temperature – SST, salinity, surface zooplankton concentration and sardine biomass in the Iberian continental shelf). The first five variables were obtained through the Copernicus Marine system with a daily temporal resolution. 8-day composites were then created using the calc function of the raster R package. The 8-day composites were extracted to the segment centroids and prediction points, according to their location and date, using the raster package. The values for the annual sardine biomass in Iberian shelf waters were retrieved from (ICES, 2024).

### 2.3 Data preparation

A dataset of common dolphin observations was created, with information concerning the number of individuals in the group, distance to the transect line and geographic location. A second dataset was created, with information regarding the distance travelled in the segment, and the geographic location and environmental conditions at the centroid. Transect segments were created, each corresponding to five minutes of sampling, resulting in 48,875 segments (mean length: 1.28 km, SD = 0.31 km). Observation and segment data were labeled to match the observations to the segment where they occurred during modeling. Geo-referenced prediction grids, with a 1 km^2^ resolution, representing the Portuguese EEZ were created using QGIS (QGIS Development Team, 2024). The shapefile used to create the grid was downloaded from (ArcGIS Hub, 2024). Spatially explicit environmental conditions were attributed to the segment dataset and prediction grids using the extract function from the raster package. Each prediction grid reflected the environmental conditions at the midpoint of each season (winter, spring, summer and autumn) throughout the sampling period (2004-2020).

### 2.4 Two-stage density surface modelling

A two-stage density surface modelling approach was used, starting with fitting detection functions, followed by fitting an environmental explicit density model, following (Miller et al., 2013).

#### 2.4.1 Detection function

Detection functions were fitted to model common dolphin’s detection probability conditional to the distance to the transect line, using the ds function from the Distance R package (Miller et al., 2019). The three key-functions (half-normal, hazard-rate and uniform) were tested, together with differing numbers of cosine adjustments – 0, 1 or 2 for detection functions with half-normal and uniform keys and 0 or 1 cosine adjustments for detection functions with the hazard-rate key. For all detection functions, the mono_method argument was “slsqp” and the optimizer was “R”. No covariates besides distance were included in the detection function. Priority to the shape of the detection process was given to choose the appropriate function to correct dolphin counts per transect segment.

#### 2.4.2 Spatial model fitting

Before modelling, correlations between environmental variables were calculated, and if the correlation was greater than 0.7 (see Figure SX), the correlated variables were not included in the same model and were tested in different models (see Table SX).

Generalized Additive Models (GAMs) were fitted using the dsm function of the dsm R package (Miller et al., 2013). These GAMs modelled common dolphin counts per transect segment, corrected for detectability, from smooth functions of the considered environmental variables. The model structure is

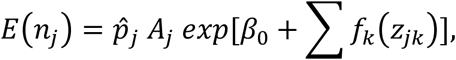

where *n*_*j*_ is the count in the *j*^*th*^ segment, *p̂*_*j*_ is the estimated detection probability in the *j*^*th*^ segment, *β*_0_ is an intercept term, the *f*_*k*_s are the smooth functions of the covariates and *z*_*jk*_ is the value of the *k*^*th*^ variable in the *j*^*th*^ segment (Miller et al., 2013).

Models were fitted using all transect segments and common dolphin sightings on effort. A stepwise backward selection was done based on AIC to select the final model. The smoothing parameter selection criterion was REML. To prevent overfitting, restrictions on the maximum degrees of freedom for all variables’ smooth functions were implemented (k = 4). As recommended in (Miller et al., 2013), the tweedie distribution was used for modelling.

### 2.5 Prediction and Variance Estimation

The final model was used to predict densities in space, using the predict function from the dsm R package. For each cell of a prediction grid, an abundance value was calculated from the environmental conditions and the cell’s area (its offset). This was done for all prediction grids. The abundance in each cell is given by

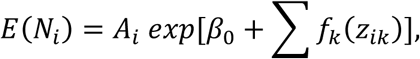

where *N*_*i*_ is the abundance in the *i*^*th*^ prediction grid cell, *β*_0_ is an intercept term, the *f*_*k*_s are the smooth functions of the covariates and *z*_*ik*_is the value of the *k*^*th*^variable in the *i*^*th*^prediction grid cell (Miller et al., 2013).

The total abundance in the study area, for each season and year, was calculated by summing the predicted abundance of all cells in the respective prediction grid. Predicted densities were the mapped in space using the ggplot2 and viridis R packages (Garnier et al., 2024; Wickham et al., 2024). To estimate confidence intervals and the coefficients of variation (CVs) associated to the abundance estimates, the dsm_var_gam function from the dsm R package was used.

## 3 RESULTS

In total, 1151 common dolphin sightings were recorded, 737 of which were during effort and used for modelling. The mean group size was 10.16 individuals (SD = 15.87) and ranged from lone animals to large aggregations of 200 dolphins. In general, common dolphins were mostly sighted in shelf waters, with greater aggregations on the northern coast (Figure 1). The environmental conditions during sampling and where common dolphins were detected are summarized in Table 2.

**TABLE 2.** Summary statistics of the environmental conditions during sampling and where common dolphin sightings on effort were recorded.

| Variable | Mean | Median | SD | Range | Temporal Resolution |
| --- | --- | --- | --- | --- | --- |
| Depth – Sampling | -670.9 | -100.9 | 1,393.0 | -5,373.3 – -0.3 | NA |
| Depth – Sightings | -271.5 | -97.2 | 632.8 | -5,122.8 – -19.2 |  |
| Slope – Sampling | 1.43 | 0.37 | 2.94 | 0.00 – 30.55 | NA |
| Slope – Sightings | 1.57 | 0.33 | 3.35 | 0.01 – 25.74 |  |
| DTC – Sampling | 42,844 | 14,588 | 78,059 | 1.4 – 370,794 | NA |
| DTC – Sightings | 23,670 | 16,705 | 36,965 | 711 – 366,450 |  |
| DT200 – Sampling | 28,108 | 14,159 | 43,121 | 3 – 322,749 | NA |
| DT200 – Sightings | 16,192 | 10,414 | 18,894 | 4 – 182,113 |  |
| DT1000 – Sightings | 35,944 | 27,805 | 36,190 | 2 – 304,563 | NA |
| DT1000 – Sightings | 25,297 | 23,415 | 18,283 | 7 – 169,332 |  |
| SST – Sampling | 16.48 | 16.11 | 2.32 | 11.14 – 26.10 | 8D |
| SST – Sightings | 15.88 | 15.65 | 2.04 | 11.99 – 21.70 |  |
| CHL - Sampling | 1.19 | 0.86 | 1.05 | 0.05 – 9.00 | 8D |
| CHL – Sightings | 1.34 | 1.06 | 1.05 | 0.08 – 5.66 |  |
| CHL-lag – Sampling | 1.15 | 0.75 | 1.06 | 0.06 – 8.25 | 8D |
| CHL-lag – Sightings | 1.20 | 0.96 | 0.93 | 0.09 – 6.04 |  |
| Salinity – Sampling | 35.49 | 35.67 | 0.93 | 18.19 – 36.81 | 8D |
| Salinity – Sightings | 35.52 | 35.62 | 0.56 | 29.31 – 36.47 |  |
| Zooplankton – Sampling | 2.91 | 2.63 | 1.90 | 0.08 – 32.03 | 8D |
| Zooplankton – Sightings | 3.43 | 3.06 | 1.97 | 0.48 – 12.62 |  |
| Biomass – Sampling | 328,417 | 289,030 | 170,357 | 116,989 – 621,202 | Y |
| Biomass – Sightings | 327,062 | 381,927 | 153,358 | 116,989 – 621,202 |  |

### 3.1 Modelling Results

The chosen detection function had a uniform key-function and two cosine adjustments (Figure 2). The estimated detection probability was 0.462 (CV = 0.041). The selected model included smooths of DTC, DT1000, SST, CHL, CHL-lag, Salinity, Zooplankton and Biomass. The model’s output suggests that common dolphin densities peak at around 10,000 m from the coast (p = 0.011), while being negatively related to DT1000, although not statistically significant (p = 0.095). Additionally, there were significant positive trends with CHL (p ≤ 0.001), and sardine Biomass (p ≤ 0.001), as well as a positive non-significant (p = 0.065) trend with salinity. Contrasting this, the model predicts significant negative relationships between dolphin densities with SST (p ≤ 0.001) and CHL-lag (p ≤ 0.001). According to the model’s output, common dolphin densities are significantly related to Zooplankton (p ≤ 0.001), with a peak of dolphin densities between 5 *g*/*m*^2^and 10 *g*/*m*^2^, with densities increasing again as Zooplankton increases beyond 20 *g*/*m*^2^, (Figure 3).

**FIGURE 2.**
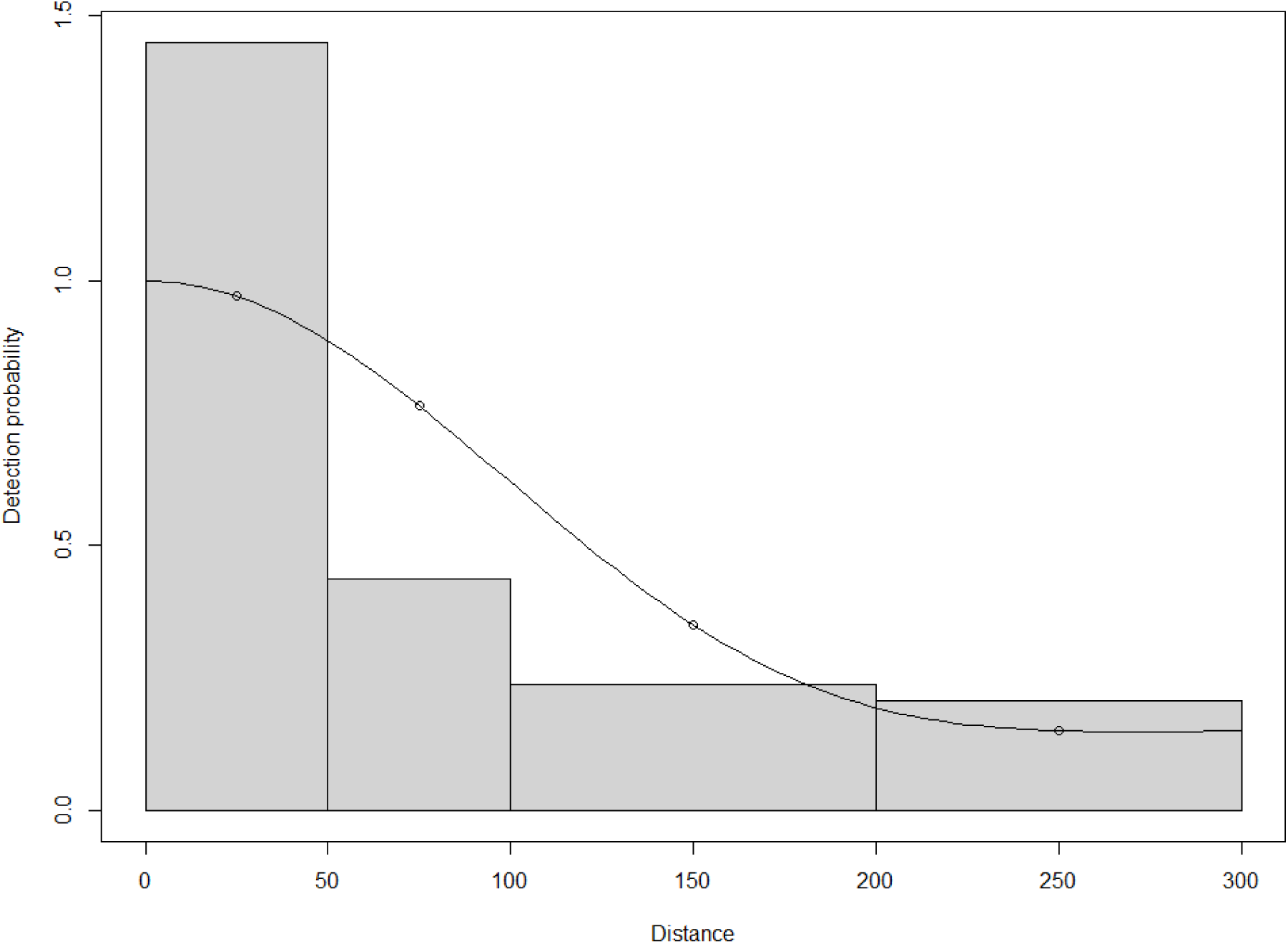
Selected detection function (line) to model common dolphin detectability according to the distance to the observer.

**FIGURE 3.**
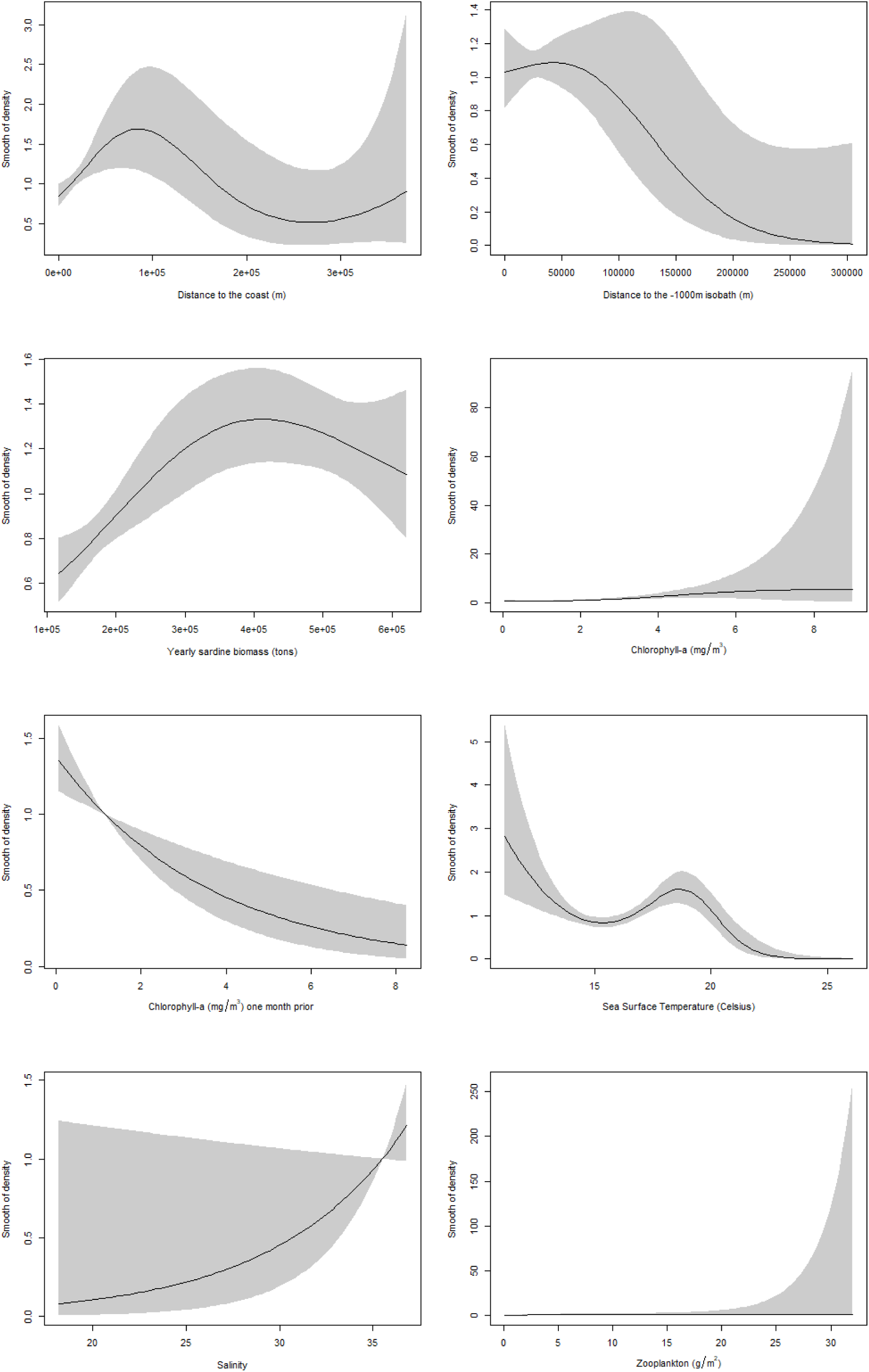
Smoothed fit plots for each covariate, showing how the estimated common dolphin density varies according to the environmental gradient. Solid lines show the mean and grey ribbons represent the standard error. See Figure S2 for smoothed fits without standard error.

### 3.2 Model Predictions

The selected model predicts both seasonal and interannual differences in the distribution and density of the common dolphin in the study area. Overall, the model predicts higher densities closer to the coast, with a patchy distribution (Figure 4). The predicted summer distribution is more concentrated in shelf waters and adjacent areas than in the remaining seasons. For them, the model still predicts higher densities close to shelf waters, however, it predicts larger numbers of common dolphins offshore than in the summer.

**FIGURE 4.**
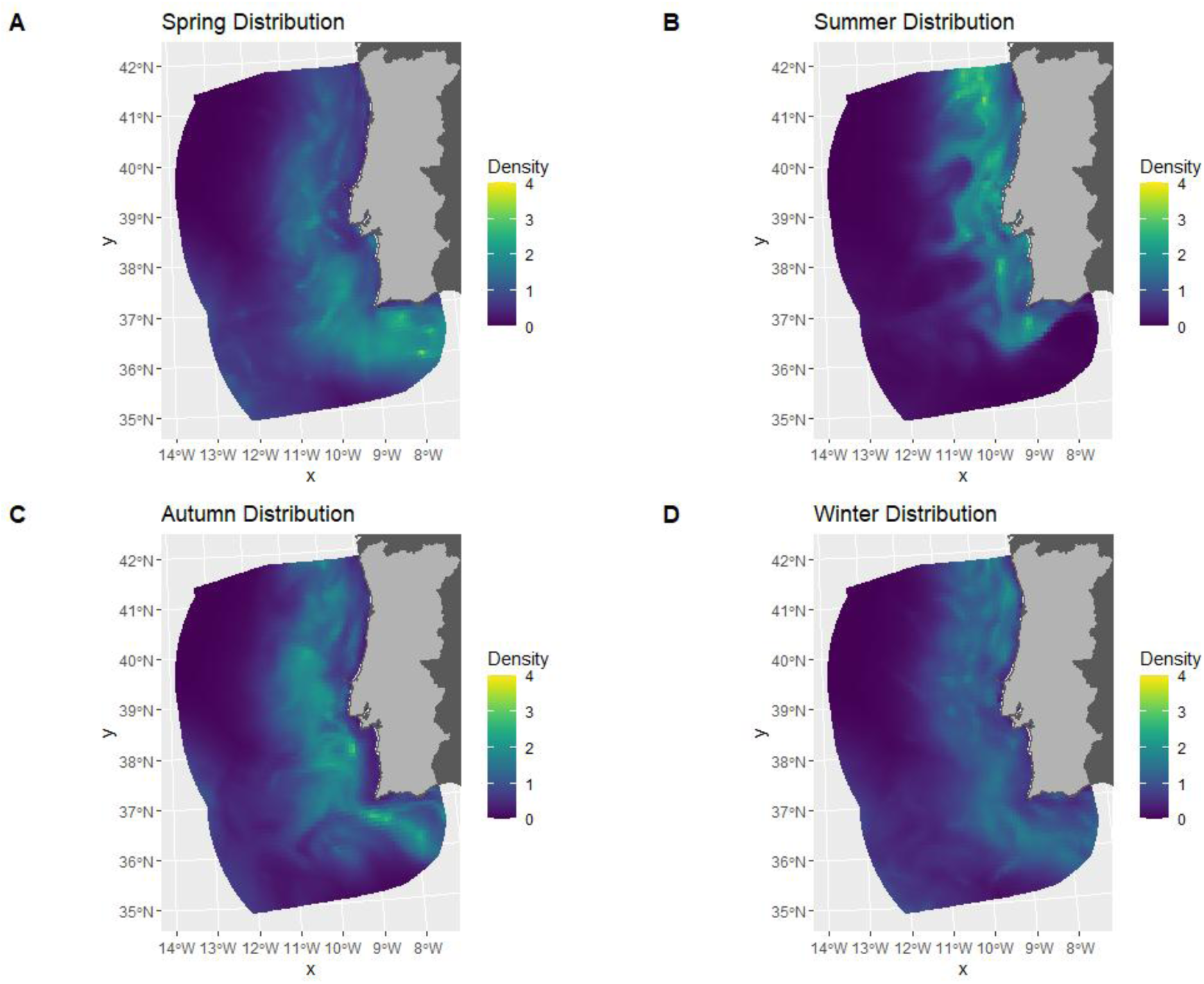
Predicted common dolphin density distribution in mainland Portugal EEZ in 2020, according to the season: A – spring, B – summer, C – autumn, D – winter. See the Supplementary Material for predictions for 2004-2019.

Generally, the predicted average density of the target species was lowest during the summer, varying from 0.201 individuals/km² (CV = 0.2657), in 2004, to 0.596 individuals/km² (CV = 0.1543), in 2007. On the other hand, spring was, in general, the season for which the model predicted highest densities, varying from 0.358 individuals/km² (CV = 0.2174), in 2012, to 0.913 individuals/km² (CV = 0.1888), in 2008. For autumn, the model predicted similar values to those in spring, although generally lower, varying from 0.296 individuals/km² (CV = 0.1914), in 2014, to 0.878 individuals/km² (CV = 0.1784), in 2008. Finally, the values for winter were, usually, intermediate between values in the summer and autumn. Estimates for the mean density in this season varied between 0.317 individuals/km² (CV = 0.2161), in 2014, and 0.702 individuals/km² (CV = 0.1831), in 2008. For detailed information on the seasonal density estimates for each year, consult Table S2, in the supplementary material.

## 4 DISCUSSION

The opportunistic dataset used in this study allowed for a unique analysis on the long-term spatiotemporal variability of common dolphin densities across such a dynamic region as the Portuguese EEZ. Nevertheless, as data collection was dedicated to seabirds and not marine mammals, some issues in detectability may arise. Most common dolphin sightings (385 out of 737) occurred in the first distance category, 0-50 m from the observer, Figure 2. This may indicate that the observers only detected common dolphins after they moved towards the vessel. If that was the case, then the density estimates would be biased high (Buckland et al., 2001). We tried to mitigate this issue by selecting the detection function based on the shape of the process instead of model fitting metrics like AIC, as using AIC would select a detection model that predicts higher detection closer to the observer, Table S3 and Figure S4, resulting in an increase in the bias. The obtained density estimates for the summer, ranging from 0.201 to 0.596 individuals/km², overlap with the estimate for the summer of 2022 obtained in (Gilles et al., 2023) of 0.363 individuals/km². Therefore, it’s likely that our choice in the detection model mitigated, at least, some impacts of the biased detectability.

In general, the common dolphin distribution predicted by the selected model was similar to what has been established in the literature (Gilles et al., 2023; Moura et al., 2012), as the smooths for DTC and DT1000, Figure 3, predict higher densities relatively close to the coast and near the -1000 m isobathymetric line, rather than offshore waters. Our predictions suggest that the species aggregates in larger numbers around shelf waters, with a patchy distribution, Figure 4, in agreement with (Moura et al., 2012). While the model still predicts the presence of common dolphins in offshore environments, predicted densities there were lower than in shelf waters, in agreement with the findings of (Gilles et al., 2023).

The model also predicts a positive relationship between common dolphin density and the sardine biomass in Iberian shelf waters, Figure 3. This variable explains most of the high interannual variation of common dolphin densities, as the years with lower density estimates (2012 – 2017, Table S2) coincided with years of lower sardine biomass in Iberian shelf waters, from 116,989 tons in 2015 to 193,646 tons in 2017, (ICES, 2024). Contrasting this, high density estimates were obtained for 2004-2008 and 2020, Table S2, which were years with high sardine biomass, varying from 381,927 tons in 2008 to 621,202 tons in 2006, (ICES, 2024). These results align with the known foraging habits of common dolphins in mainland Portugal, as its preferred prey item is the sardine, being its main food-source even when stocks were declining (Marçalo et al., 2018). While this variable seems to have a significant effect on common dolphin densities, it’s not spatially explicit. As of the writing of this paper, there aren’t any available spatially explicit products of sardine biomass in Iberian waters, which creates a limitation in our understanding of how it impacts the distribution of its predators, particularly, the common dolphin. If such products existed, and we were able to model common dolphin densities in relation to sardine biomass distribution, our density and distribution predictions would be more ecologically sensible.

The other dynamic variables (SST, CHL, CHL-lag, Salinity and Zooplankton) were used for modelling as proxies for prey distribution. The model predicts higher common dolphin densities in colder waters (below 20° C) with high chlorophyll-a and zooplankton concentrations, Figure 3. This concurs with the species distribution patterns. Common dolphins in the Eastern Tropical Pacific and in the Eastern North Atlantic have been found to be associated with colder waters, rich in nutrients (Au & Perrymanl, 1985; Fernandez et al., 2021), and the same seems to apply here. Interestingly, our model results show a positive relationship between common dolphin density and chlorophyll-a concentration in the week of the sighting, but a negative relationship with a chlorophyll-a concentration with a one-month time lag, Figure 3. This suggests that common dolphins aggregate in high primary productivity events when they are happening. Additionally, the positive relationship between common dolphin densities and salinity, combined with a preference for colder waters, matches the habitat preference of the sardine, which has been found to prefer a water temperature below 20° C and high sea surface salinity (Lima et al., 2022).

The results from the sardine biomass and the dynamic variables support that common dolphin densities are likely dependent on prey distribution. A possible explanation for the high variability of density estimates along the years is that when sardine is less abundant in mainland Portugal, common dolphins move to adjacent waters in search of prey. The same line of thought can be applied for the seasonal densities. It’s likely that dolphins move in and out mainland Portugal EEZ seasonally and occupy different habitats in response to their prey’s movements.

In a climate change scenario, where SST is expected to increase on average from 1.2° C to 3.2° C, depending on the scenario, (Genner et al., 2017), the colder water dependent sardines are expected to have a northward shift in distribution (Lima et al., 2022). This would result in lower prey availability for mainland Portugal’s common dolphins, which preferably target sardines (Marçalo et al., 2018). Climate change impacts may be exacerbated by human activity, especially overfishing, reducing even further the sardine biomass, leading to the potential reduction of common dolphin densities in the region. However, this doesn’t account for the dolphin’s adaptability. Cetaceans, such as common dolphins are highly adaptable, and common dolphins have been found to shift their main prey items in Galician waters, in proximity to the study area, in response to prey availability (Santos et al., 2013). Nevertheless, this is contrary to what has been reported for mainland Portugal, (Marçalo et al., 2018), and our results also support a link between common dolphin densities and sardine biomass. Therefore, we consider that sardine stock management is likely going to be important for the conservation of the common dolphin in mainland Portugal.

Despite the opportunistic nature of the dataset, the relationships we established between the considered environmental variables with common dolphin densities agree with the available literature on the species ecology in the study area and proximal regions (Fernandez et al., 2021; Gilles et al., 2023; Marçalo et al., 2018; Moura et al., 2012). Moreover, we were able to establish a relationship between the common dolphin abundance and its preferred prey item, the sardine. In this sense, we were able to show how common dolphin densities and distribution change interannually and seasonally, adding new information on the species ecology, especially in offshore environments and during winter, where and when survey effort is typically lower (Gilles et al., 2023; Virgili et al., 2024). However, the detectability issues in this dataset can’t be disregarded, despite our attempt to mitigate them. It may be interesting to try and integrate this dataset with other datasets from dedicated marine mammal transects. That would bring data with less detectability issues, while making use of the new important information this long-term dataset provides.

## Notes

### Competing Interest Statement

The authors have declared no competing interest.

### Funding

- CEAUL

