## Supplementary_material for "Seasonal and interannual variation of common dolphin’s densities in Portuguese waters"

**TABLE S1 –** Summary of Generalized Additive Models considered in this study. [*] indicates the selected model.

| **Explanatory variables** | **Deviance Explained** | **AIC** | **Δ** **AIC** |
| --- | --- | --- | --- |
| s(Slope) + s(DTC) + s(DT1000) + s(SST) + s(CHL) + s(CHL-lag) + s(Salinity) + s(Zooplankton) + s(Biomass) | 5.69% | 11302.18 | 2.29 |
| s(DTC) + s(DT1000) + s(SST) + s(CHL) + s(CHL-lag) + s(Salinity) + s(Zooplankton) + s(Biomass) [*] | 5.66% | 1299.89 | 0.00 |
| s(DTC) + s(DT1000) + s(SST) + s(CHL) + s(CHL-lag) + s(Zooplankton) + s(Biomass) | 5.49% | 11302.10 | 2.22 |
| s(DTC) + s(SST) + s(CHL) + s(CHL-lag) + s(Salinity) + s(Zooplankton) + s(Biomass) | 5.45% | 11301.08 | 1.19 |
| s(Slope) + s(Depth) + s(DT1000) + s(SST) + s(CHL) + s(CHL-lag) + s(Salinity) + s(Zooplankton) + s(Biomass) | 5.33% | 11309.39 | 9.50 |
| s(Depth) + s(DT1000) + s(SST) + s(CHL) + s(CHL-lag) + s(Salinity) + s(Zooplankton) + s(Biomass) | 5.29% | 11307.96 | 8.08 |
| s(DT1000) + s(SST) + s(CHL) + s(CHL-lag) + s(Salinity) + s(Zooplankton) + s(Biomass) | 5.21% | 11305.55 | 5.67 |
| s(Depth) + s(SST) + s(CHL) + s(CHL-lag) + s(Salinity) + s(Zooplankton) + s(Biomass) | 4.76% | 11313.78 | 13.90 |
| s(Slope) + s(DT200) + s(SST) + s(CHL) + s(CHL-lag) + s(Salinity) + s(Zooplankton) + s(Biomass) | 4.99% | 11312.86 | 12.98 |
| s(DT200) + s(SST) + s(CHL) + s(CHL-lag) + s(Salinity) + s(Zooplankton) + s(Biomass) | 4.99% | 11310.89 | 11.01 |
| s(DT200) + s(SST) + s(CHL) + s(CHL-lag) + s(Zooplankton) + s(Biomass) | 4.83% | 11312.92 | 13.03 |

**TABLE S2 –** Summary of model-based common dolphin density predictions for mainland Portugal EEZ, per year and season. CI stands for Confidence Interval and CV for Coefficient of Variation.

| **Year** | **Season** | **Mean Density**  **(**$\boldsymbol{n/k}\boldsymbol{m}^{\boldsymbol{2}}$**)** | **Abundance**  **(point estimate)** | **95% CI** | **CV** |
| --- | --- | --- | --- | --- | --- |
| 2004 | Spring  Summer  Autumn  Winter | 0.751  0.201  0.754  0.632 | 236,311  63,140  237,213  198,709 | (165,920; 336,564)  (37,841; 105,354)  (166,758; 337;436)  (138,301; 285,501) | 0.1819  0.2657  0.1813  0.1865 |
| 2005 | Spring  Summer  Autumn  Winter | 0.826  0.460  0.702  0.632 | 259,902  144,671  220,805  198,765 | (178,151; 379,167)  (105,871; 197,690)  (159,382; 305,900)  (138,006; 286,274) | 0.1945  0.1603  0.1675  0.1878 |
| 2006 | Spring  Summer  Autumn  Winter | 0.655  0.304  0.505  0.522 | 206,207  95,602  158,929  164,171 | (131,807; 322,602)  (64,888; 140,855)  (107,448; 235,076)  (103,189; 261,191) | 0.2313  0.1997  0.2017  0.2403 |
| 2007 | Spring  Summer  Autumn  Winter | 0.681  0.596  0.763  0.606 | 214,106  187,445  240,067  190,493 | (151,287; 303,009)  (138,771; 253,191)  (176,081; 327,305)  (132,486; 190,493) | 0.1786  0.1543  0.1591  0.1869 |
| 2008 | Spring  Summer  Autumn  Winter | 0.913  0.470  0.878  0.702 | 287,089  147,701  276,346  220,855 | (198,918; 414,340)  (102,121; 213,624)  (195,329; 390,966)  (154,716; 315,266) | 0.1888  0.1900  0.1784  0.1831 |
| 2009 | Spring  Summer  Autumn  Winter | 0.646  0.501  0.601  0.588 | 203,109  157,511  188,917  185,029 | (137,537; 299,942)  (112,874; 219,799)  (132,931; 268,482)  (124,249; 275,543) | 0.2009  0.1713  0.1808  0.2053 |
| 2010 | Spring  Summer  Autumn  Winter | 0.641  0.455  0.631  0.520 | 201,782  143,161  198,369  163,664 | (139,247; 292,401)  (103,922; 197,214)  (139,939; 281,196)  (112,083; 238,983) | 0.1910  0.1645  0.1794  0.1950 |
| 2011 | Spring  Summer  Autumn  Winter | 0.576  0.470  0.537  0.392 | 181,124  147,923  168,902  123,409 | (126,632; 259,064)  (107,582; 203,390)  (117,684; 242,412)  (85,234; 178,684) | 0.1841  0.1635  0.1859  0.1905 |
| 2012 | Spring  Summer  Autumn  Winter | 0.358  0.298  0.389  0.318 | 112,597  93,731  122,356  99,888 | (73,898; 171,563)  (65,479; 134,173)  (83,400; 179,507)  (66,269; 150,563) | 0.2174  0.1846  0.1974  0.2117 |
| 2013 | Spring  Summer  Autumn  Winter | 0.442  0.245  0.365  0.329 | 139,013  77,008  114,771  103,657 | (93,817; 205,981)  (52,352; 113,276)  (77,489; 169,990)  (67,834; 158,400) | 0.2027  0.1988  0.2024  0.2189 |
| 2014 | Spring  Summer  Autumn  Winter | 0.397  0.312  0.296  0.317 | 124,904  98,228  93,272  99,758 | (83,323; 187,237)  (67,918; 142,064)  (64,315; 135,265)  (65,626; 151,643) | 0.2088  0.1899  0.1914  0.2161 |
| 2015 | Spring  Summer  Autumn  Winter | 0.381  0.281  0.345  0.327 | 119,751  88,265  108,418  103,000 | (77,423; 185,219)  (61,432; 126,819)  (73,081; 160,842)  (66,923; 158,525) | 0.2253  0.1865  0.2033  0.2227 |
| 2016 | Spring  Summer  Autumn  Winter | 0.403  0.332  0.388  0.372 | 126,743  104,296  121,920  117,069 | (86,651; 185,386)  (75,429; 144,210)  (85,301; 174,255)  (78,548; 174,483) | 0.1959  0.1665  0.1837  0.2057 |
| 2017 | Spring  Summer  Autumn  Winter | 0.569  0.393  0.481  0.423 | 178,959  123,612  151,236  133,114 | (124,345; 257,560)  (90,085; 169,616)  (107,617; 212,534)  (91,267; 194,149) | 0.1874  0.1625  0.1749  0.1944 |
| 2018 | Spring  Summer  Autumn  Winter | 0.497  0.345  0.523  0.443 | 156,268  108,616  164,533  139,239 | (109,558; 222,894)  (77,221; 152,774)  (117,775; 229,853)  (97,941; 197,951) | 0.1827  0.1754  0.1718  0.1810 |
| 2019 | Spring  Summer  Autumn  Winter | 0.522  0.408  0.520  0.495 | 164,122  128,327  163,707  155,615 | (113,920; 236,447)  (92,781; 177,492)  (116,121; 230,795)  (108,454; 223,285) | 0.1879  0.1666  0.1766  0.1858 |
| 2020 | Spring  Summer  Autumn  Winter | 0.754  0.564  0.734  0.644 | 237,176  177,424  230,861  202542 | (162,788; 345,556)  (129,870; 242,391)  (164,792; 323,419)  (140,585; 291,805) | 0.1938  0.1602  0.1733  0.1879 |

**TABLE S3 –** Summary of detection functions tested for modelling common dolphin detectability according to the distance to the observer. [*] indicates the selected function.

| **Key function** | **N of cosine adjustments** | **Average p** | **CV** | **AIC** |
| --- | --- | --- | --- | --- |
| Half-Normal | 0 | 05008 | 0.02517 | 1998.288 |
| Half-Normal | 1 | 0.6787 | 0.05197 | 2077.666 |
| Half-Normal | 2 | 0.3334 | 0.05609 | 1836.717 |
| Uniform | 0 | 1 | 0 | 2313.888 |
| Uniform | 1 | 0.5589 | 0.02026 | 2014.100 |
| Uniform [*] | 2 | 0.4626 | 0.04111 | 1933.177 |
| Hazard-rate | 0 | 0.1674 | 0.32210 | 1807.777 |
| Hazard-rate | 1 | 0.2083 | 0.26825 | 1804.114 |

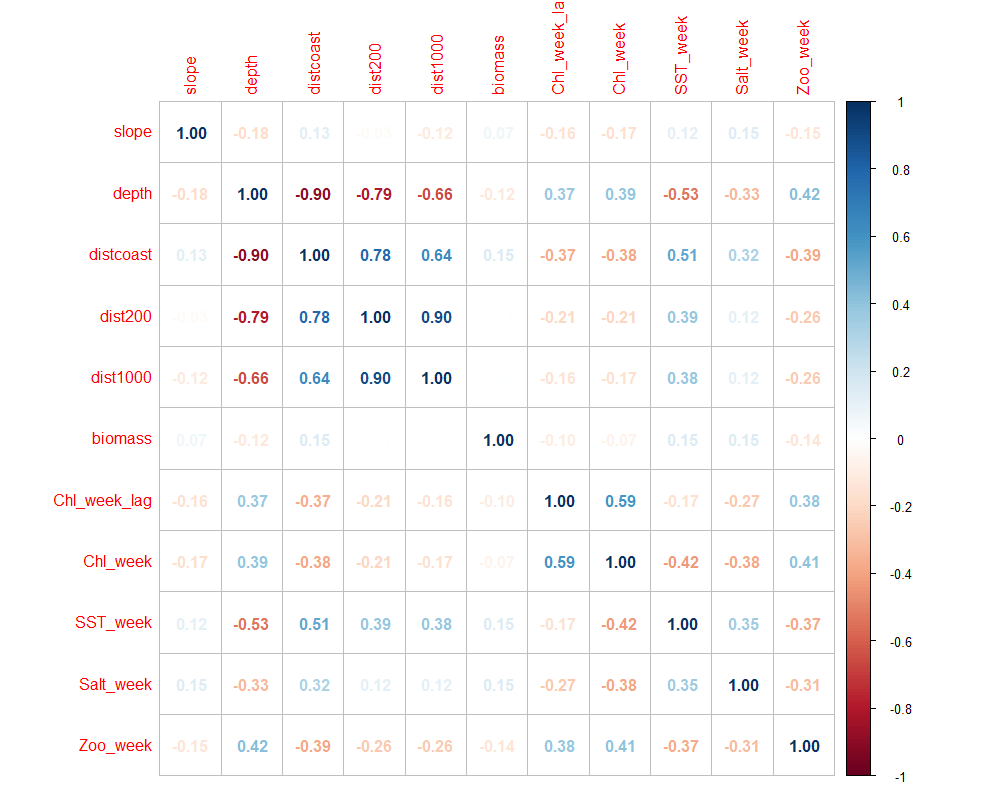

**FIGURE S1** – Correlation matrix of variables considered for modelling common dolphin densities.

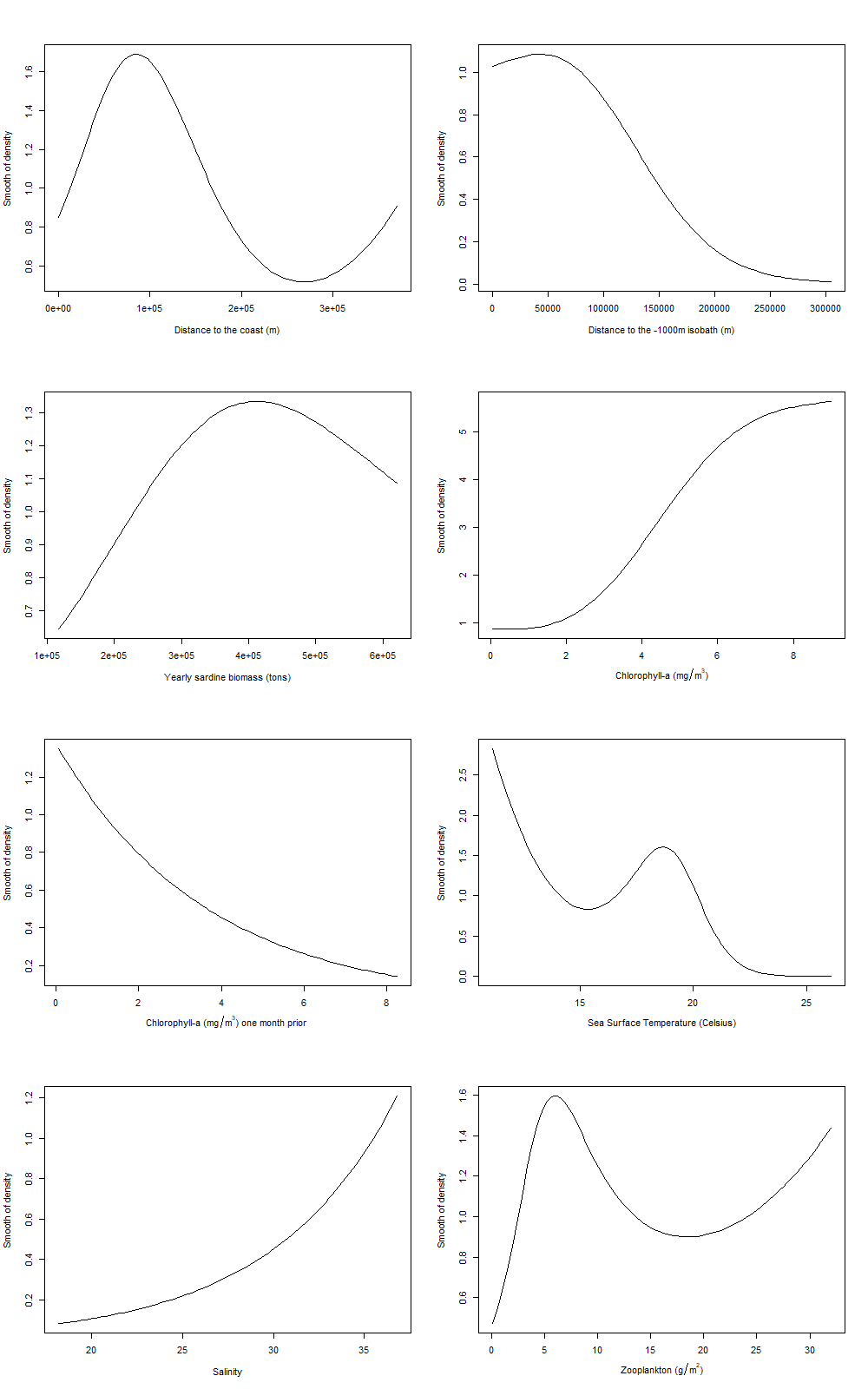

**FIGURE S2** – Smoothed fits of predictors, showing how the estimated common dolphin density varies according the environmental gradient.

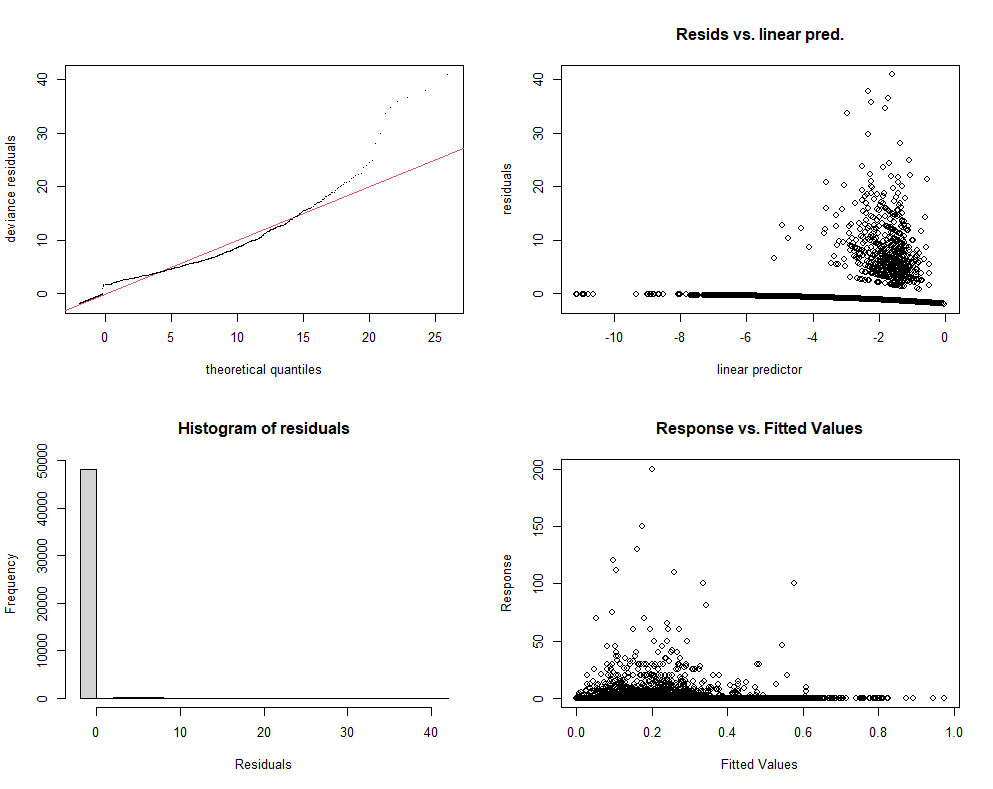

**FIGURE S3** – Diagnostic plots of the selected model.

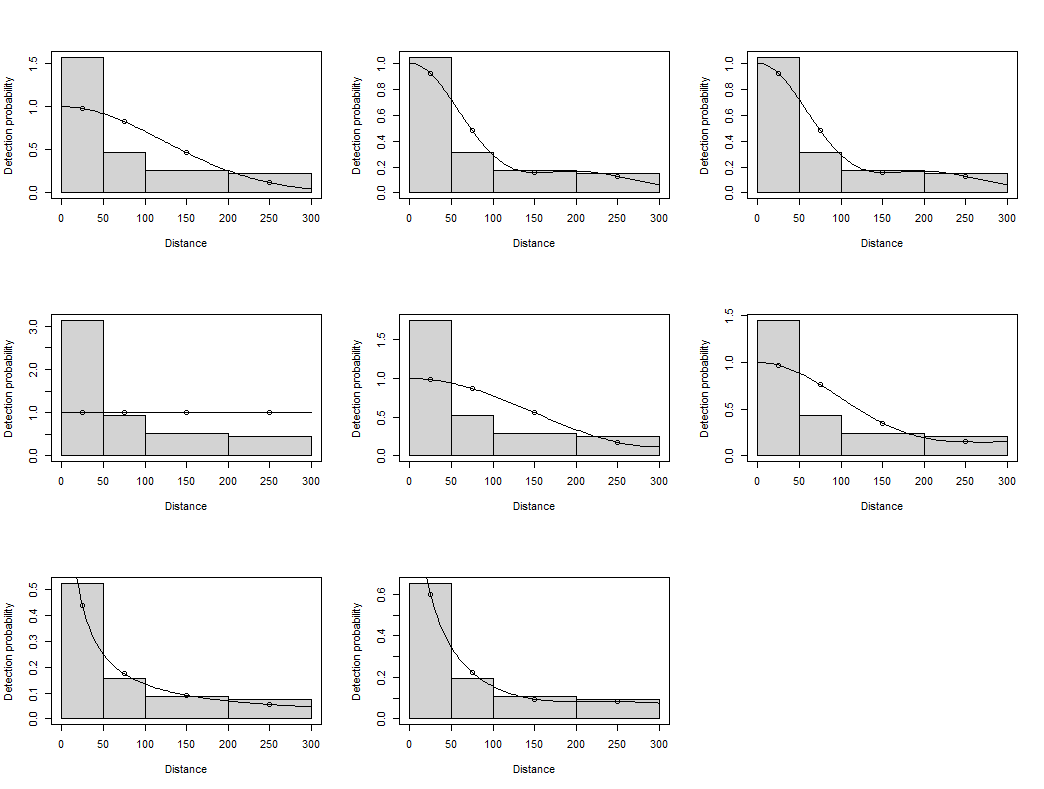

**FIGURE S4** – Plots of tested detection functions. From top to bottom, left to right: half-normal key with 0, 1 and 2 cosine adjustments, respectively; uniform key with 0, 1 and 2 cosine adjustments, respectively; hazard-rate key with 0 and 1 cosine adjustments, respectively.

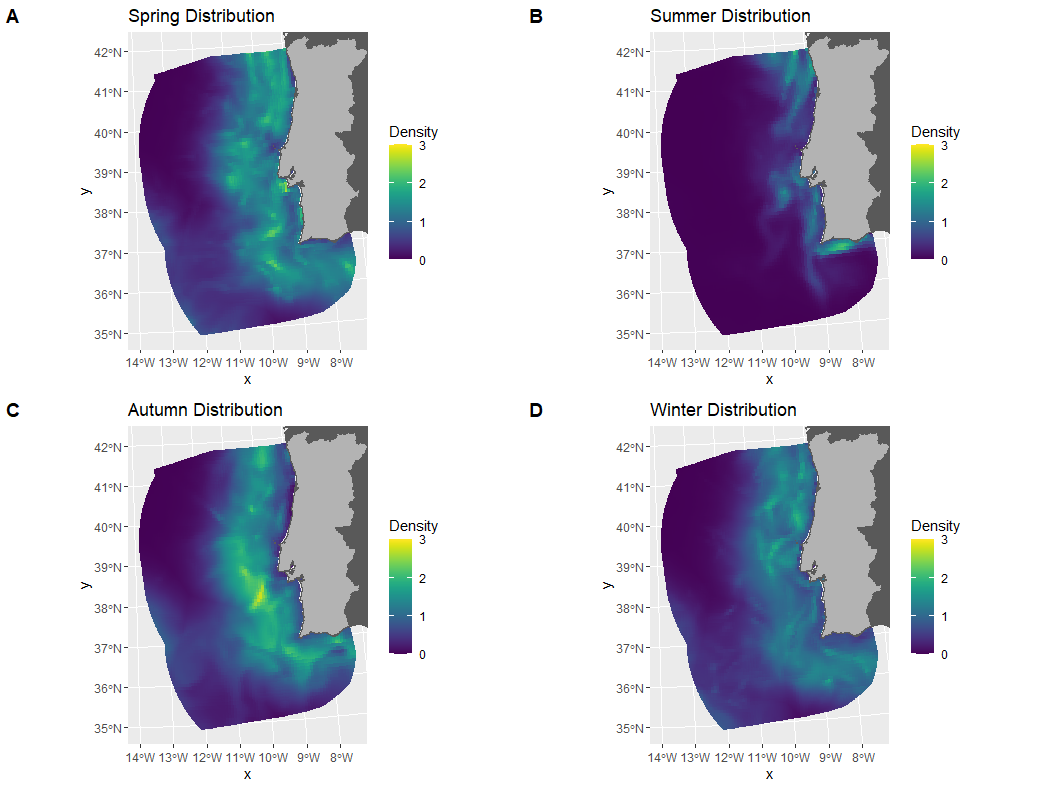

**FIGURE S5 -** Predicted common dolphin density distribution in mainland Portugal EEZ in 2004, according to the season.

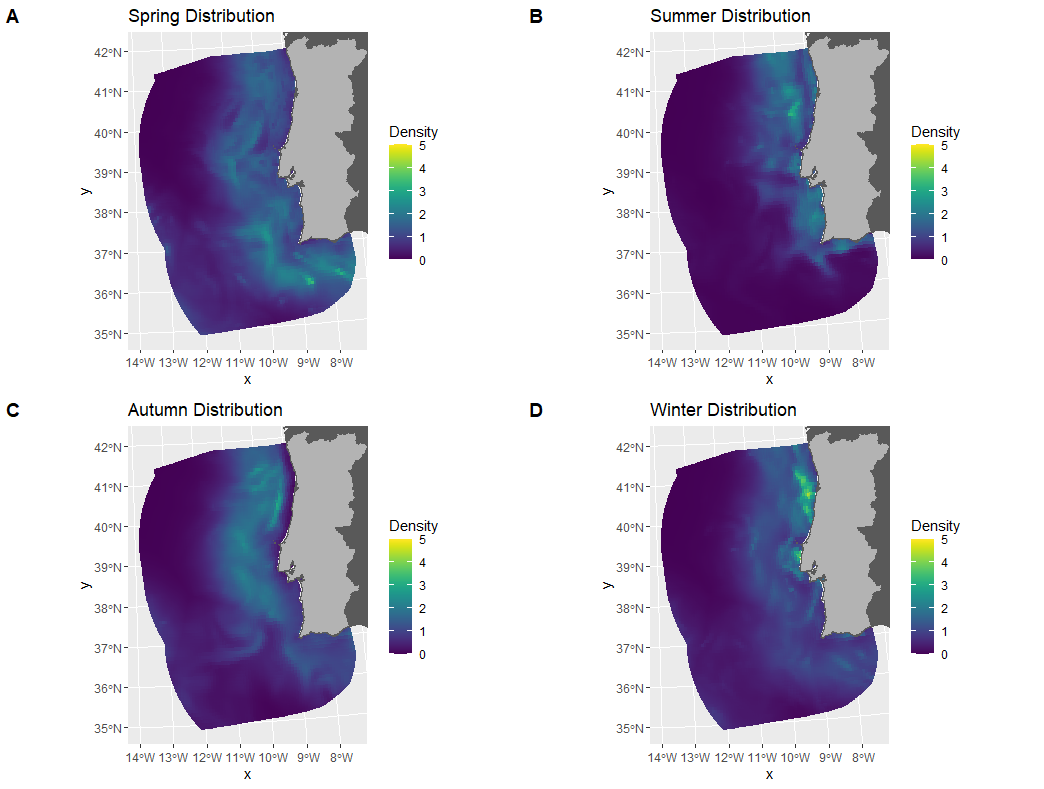

**FIGURE S6 -** Predicted common dolphin density distribution in mainland Portugal EEZ in 2005, according to the season.

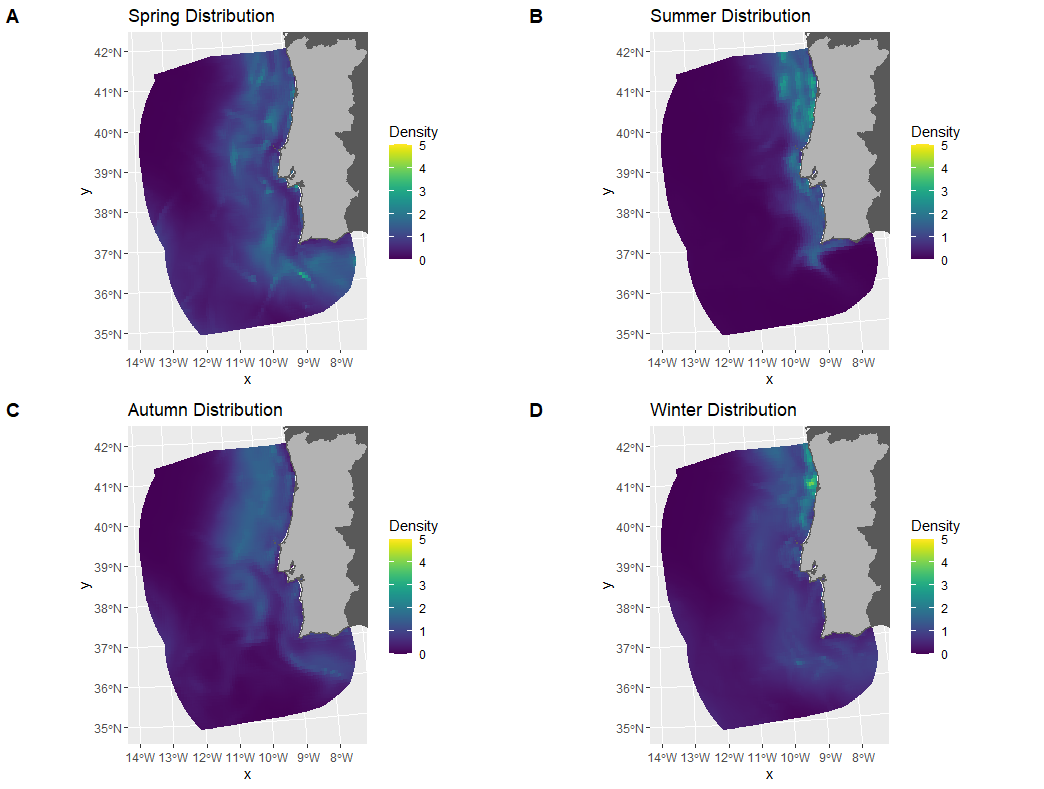

**FIGURE S7 -** Predicted common dolphin density distribution in mainland Portugal EEZ in 2006, according to the season.

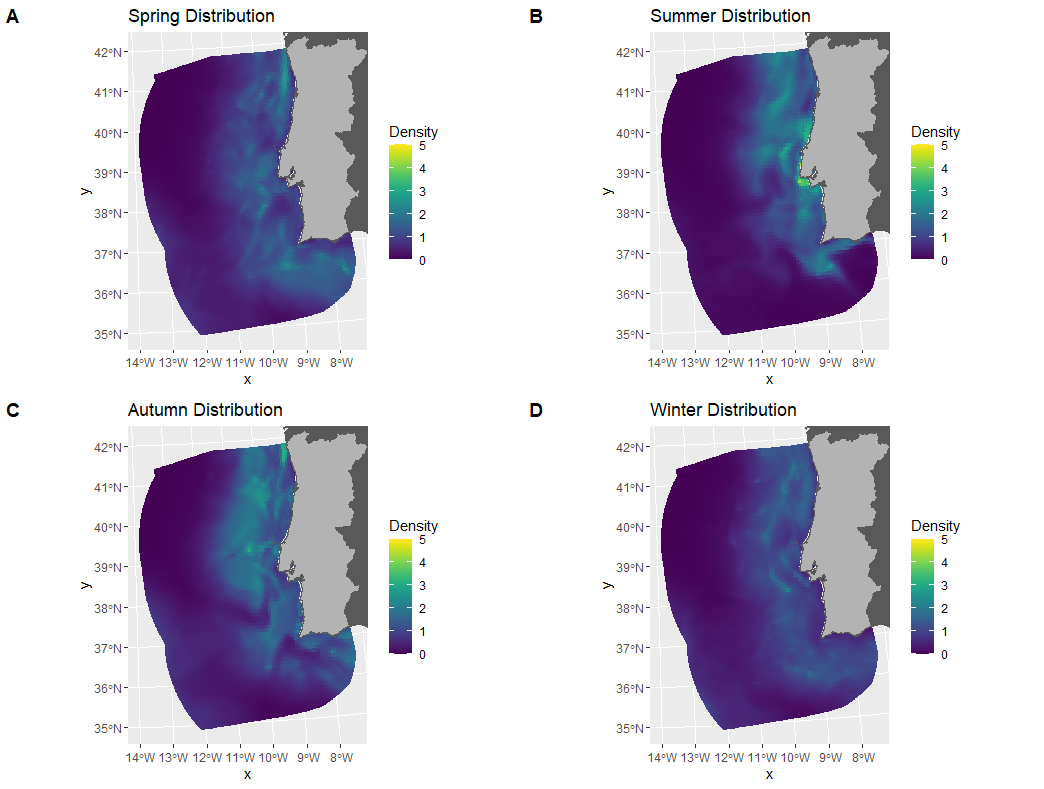

**FIGURE S8 -** Predicted common dolphin density distribution in mainland Portugal EEZ in 2007, according to the season.

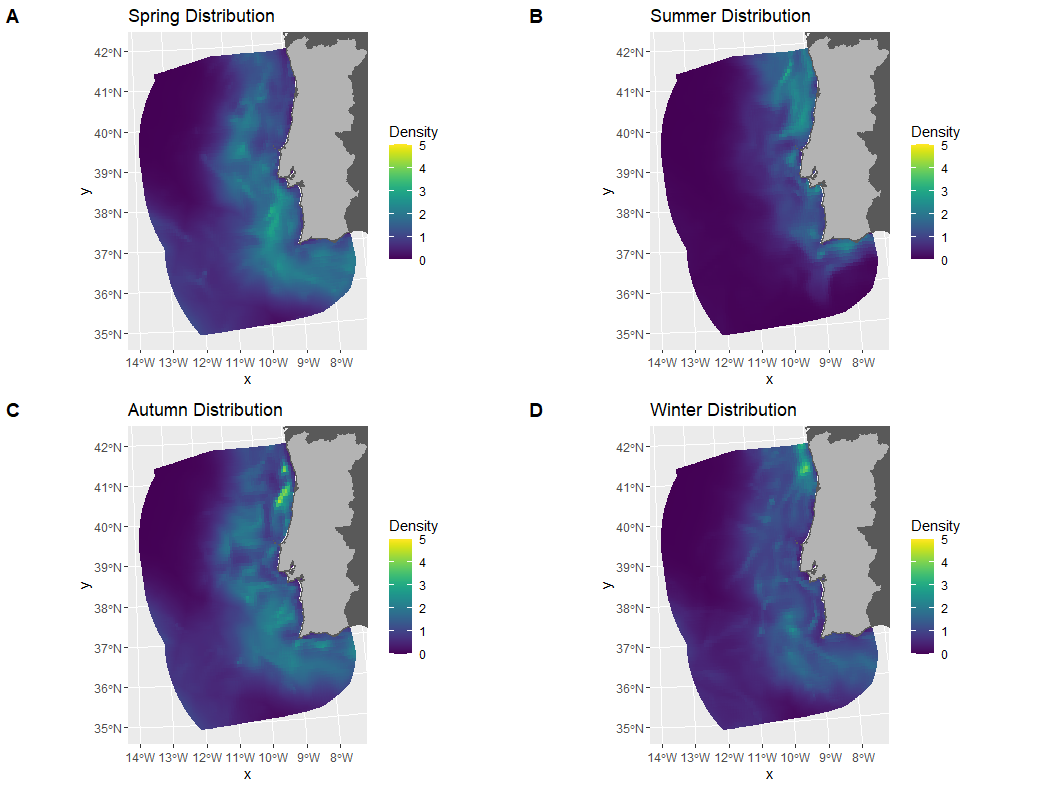

**FIGURE S9 -** Predicted common dolphin density distribution in mainland Portugal EEZ in 2008, according to the season.

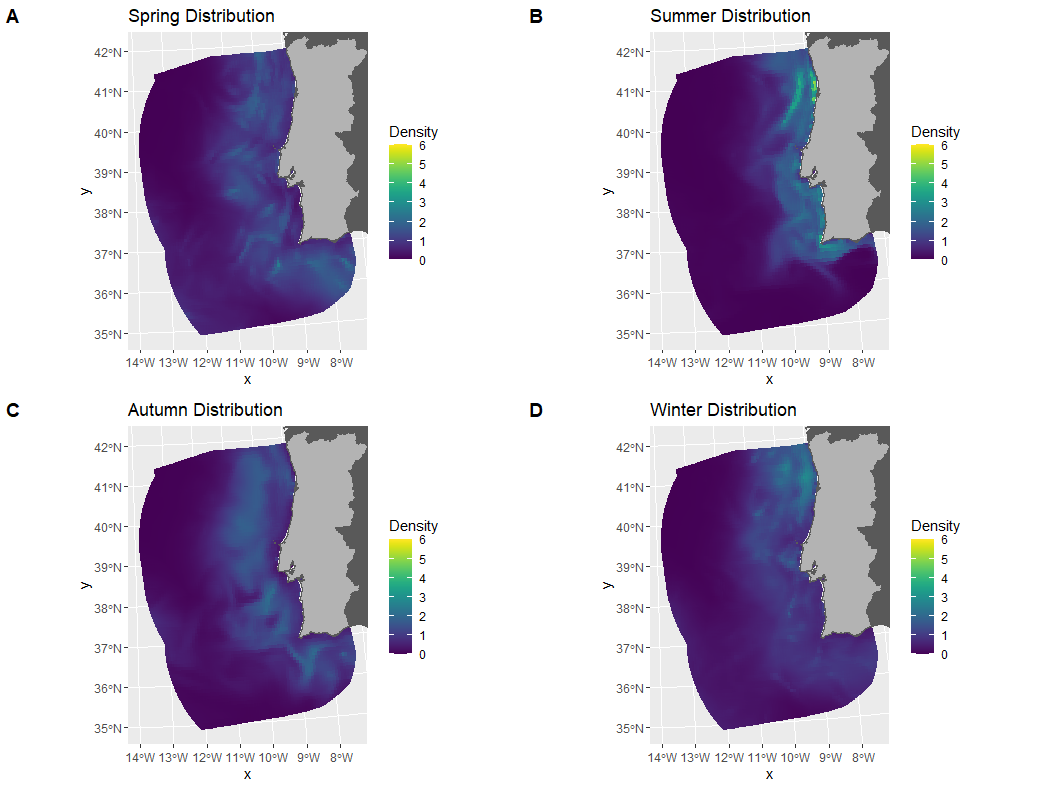

**FIGURE S10 -** Predicted common dolphin density distribution in mainland Portugal EEZ in 2009, according to the season.

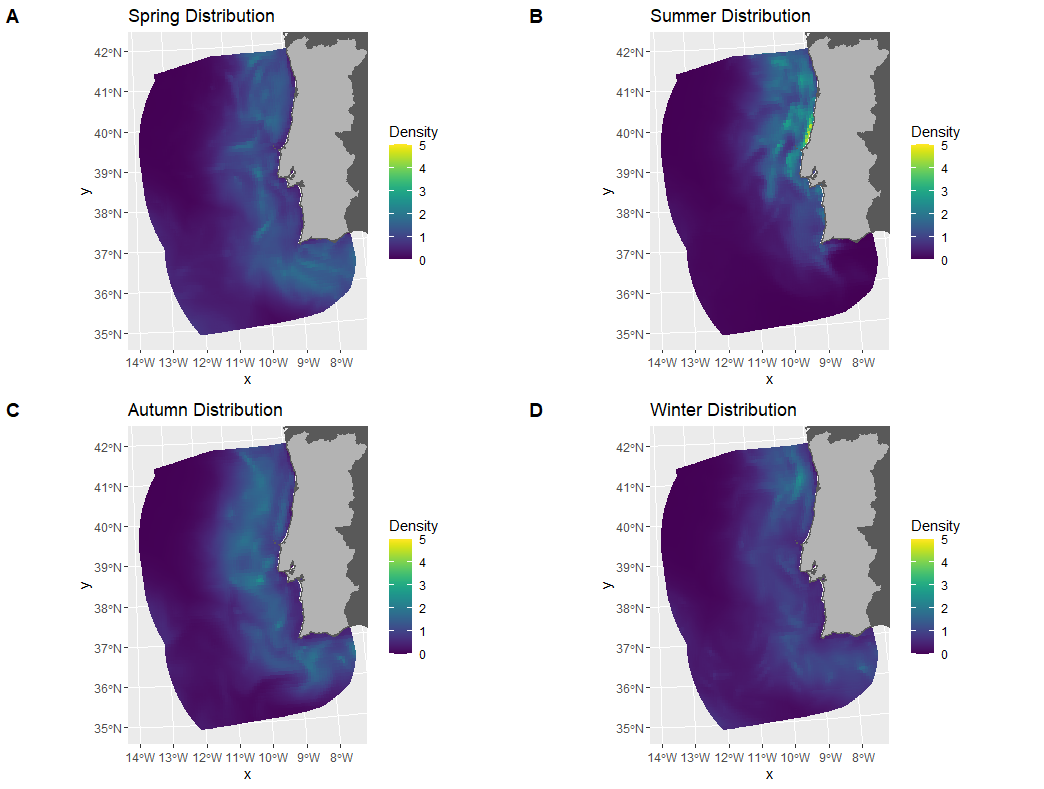

**FIGURE S11 -** Predicted common dolphin density distribution in mainland Portugal EEZ in 2010, according to the season.

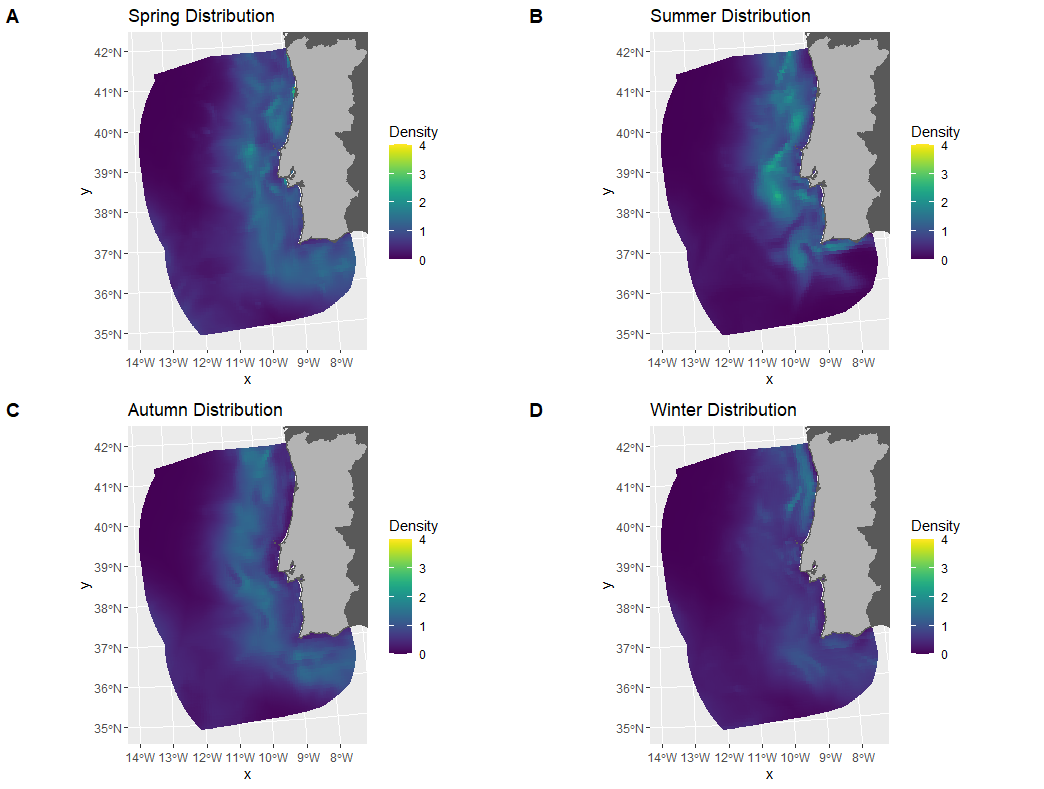

**FIGURE S12 -** Predicted common dolphin density distribution in mainland Portugal EEZ in 2011, according to the season.

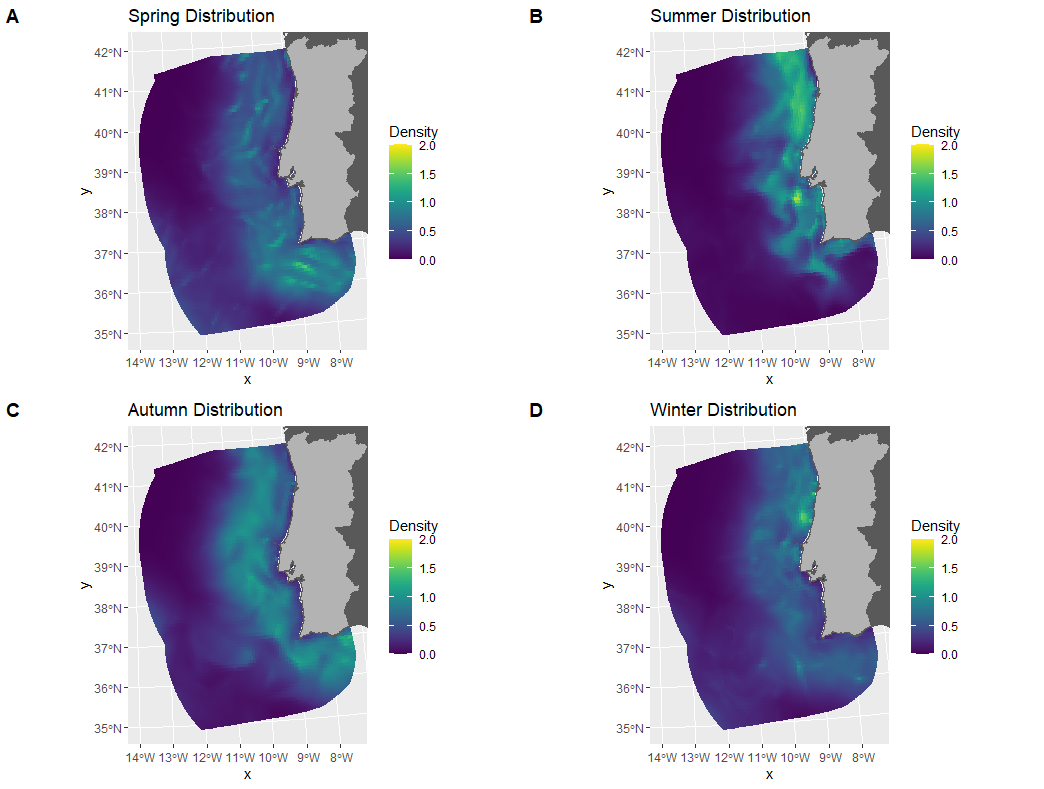

**FIGURE S13 -** Predicted common dolphin density distribution in mainland Portugal EEZ in 2012, according to the season.

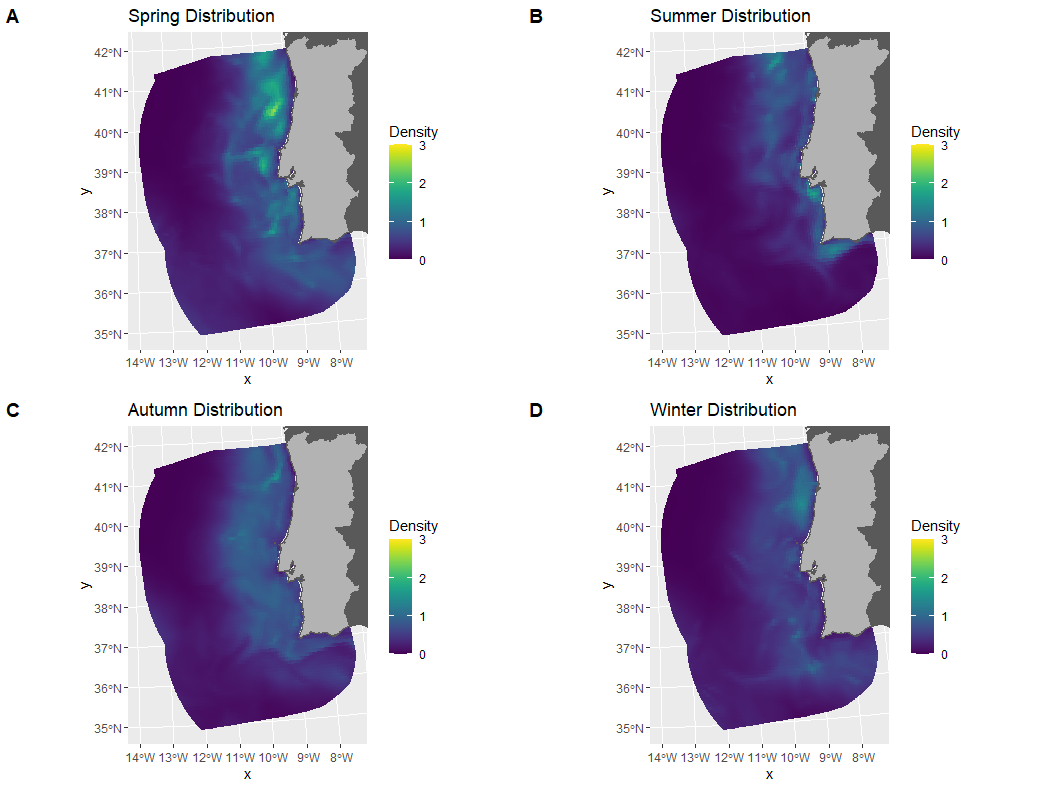

**FIGURE S14 -** Predicted common dolphin density distribution in mainland Portugal EEZ in 2013, according to the season.

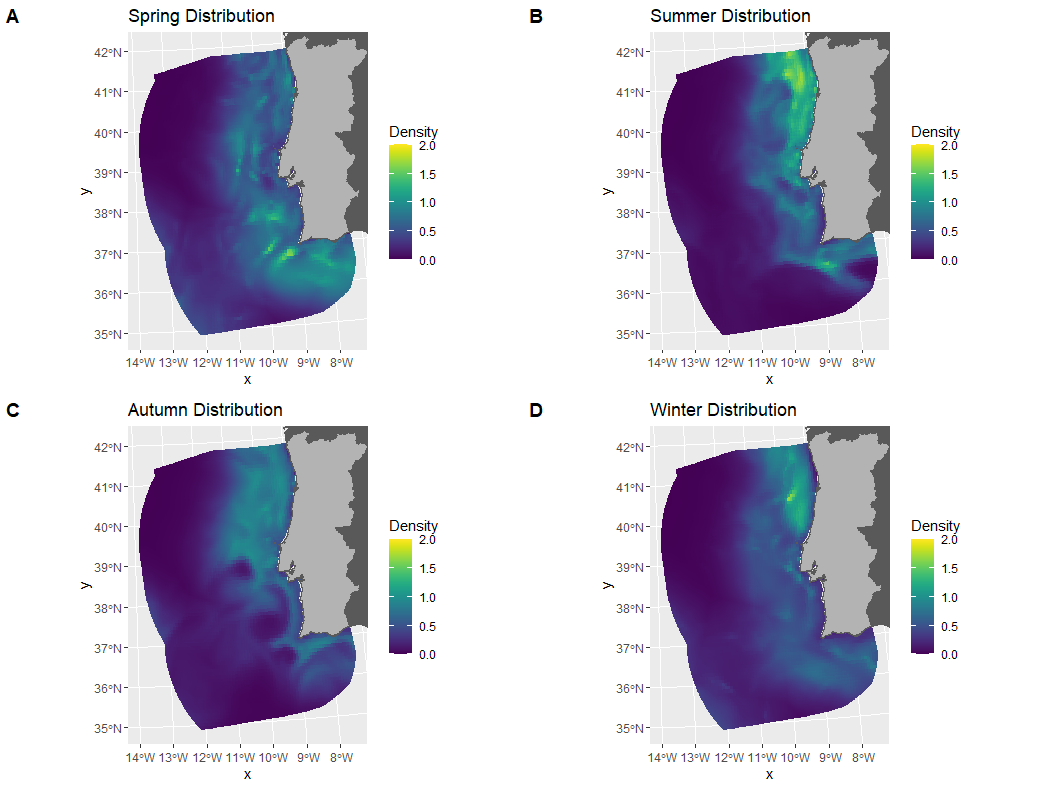

**FIGURE S15 -** Predicted common dolphin density distribution in mainland Portugal EEZ in 2014, according to the season.

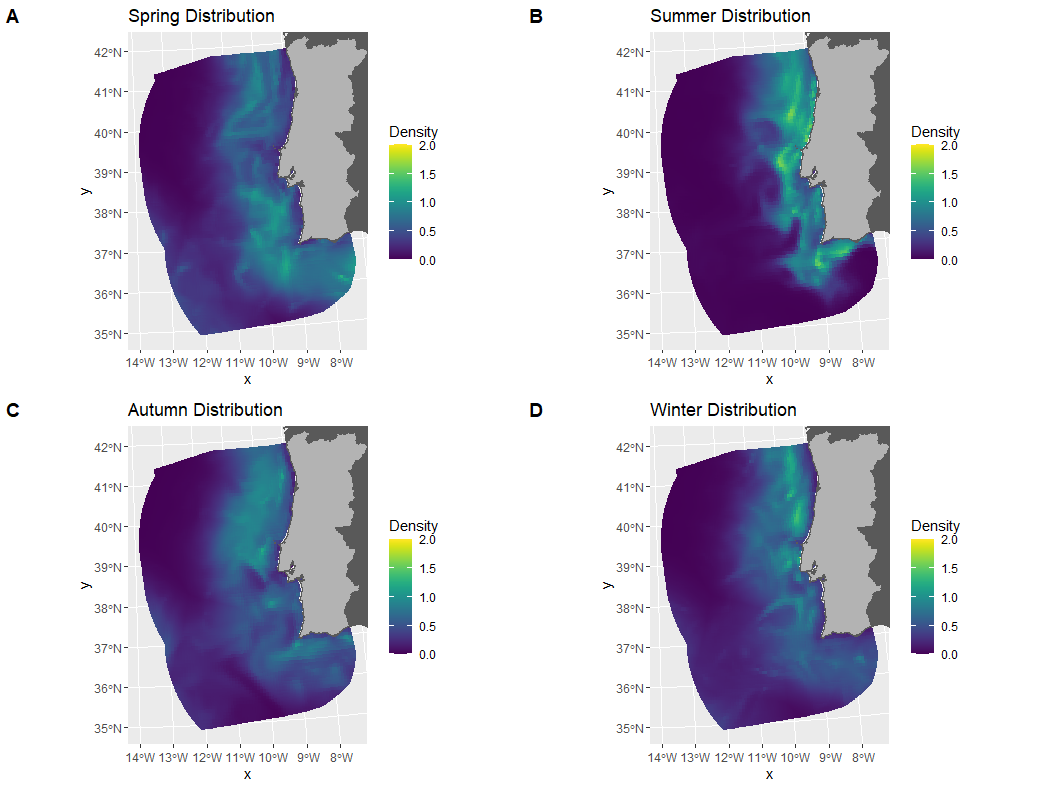

**FIGURE S16 -** Predicted common dolphin density distribution in mainland Portugal EEZ in 2015, according to the season.

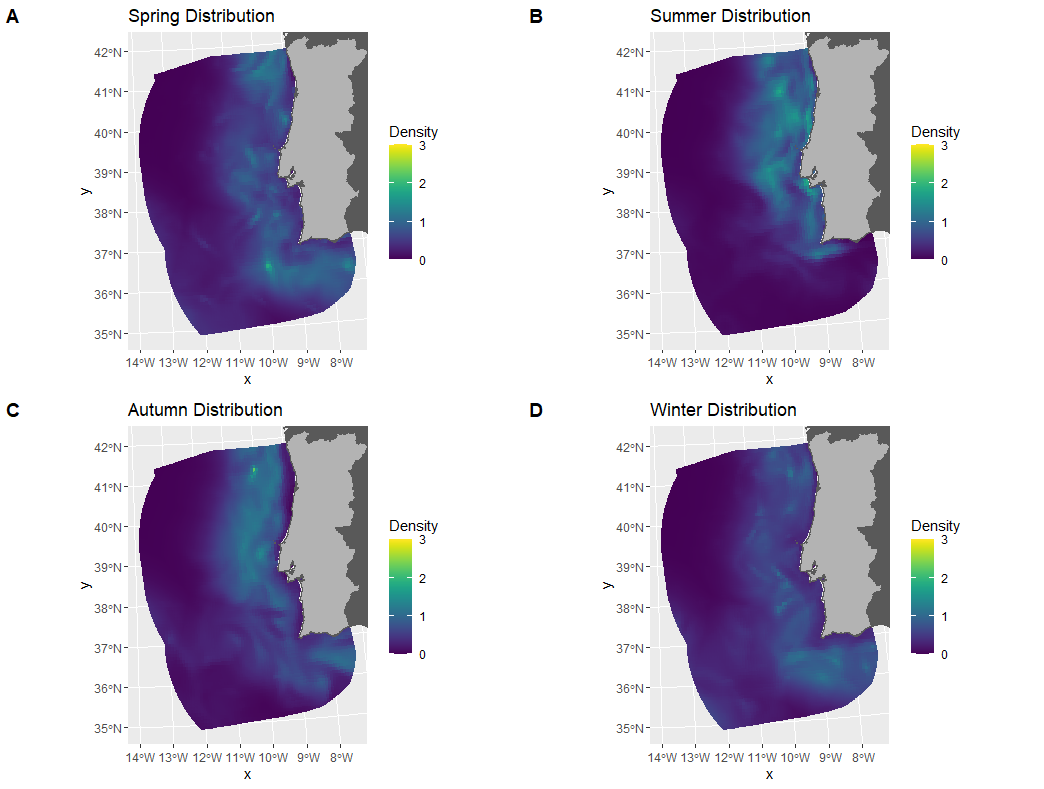

**FIGURE S17 -** Predicted common dolphin density distribution in mainland Portugal EEZ in 2016, according to the season.

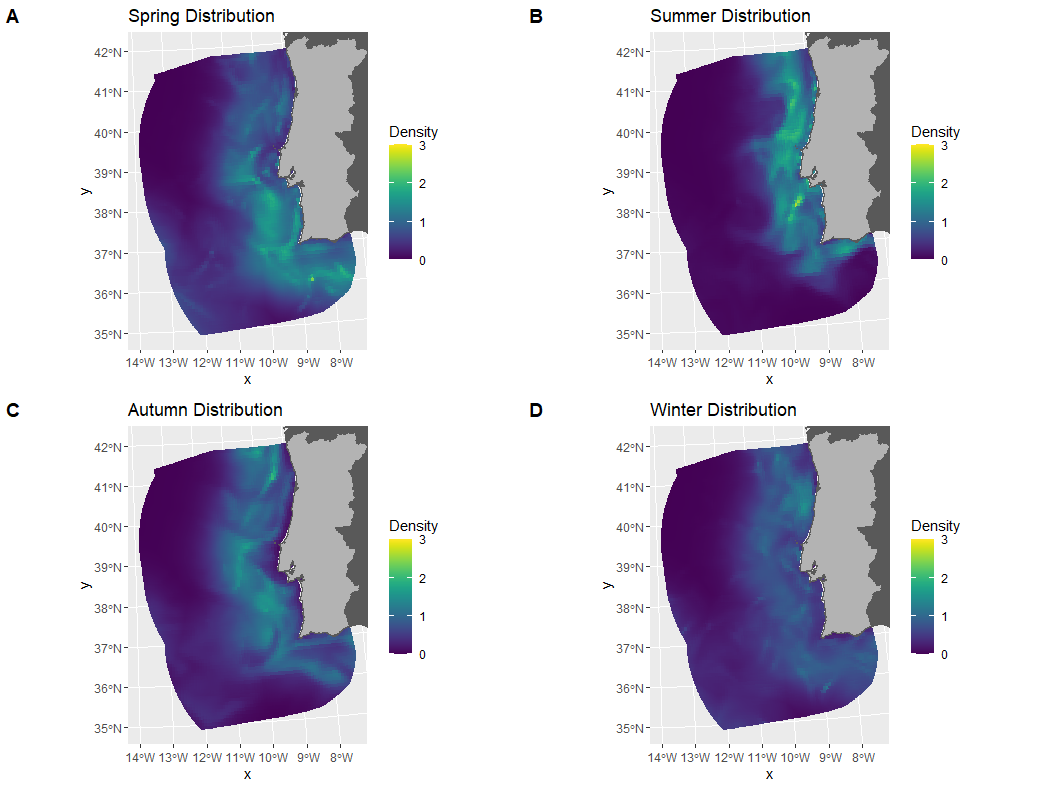

**FIGURE S18 -** Predicted common dolphin density distribution in mainland Portugal EEZ in 2017, according to the season.

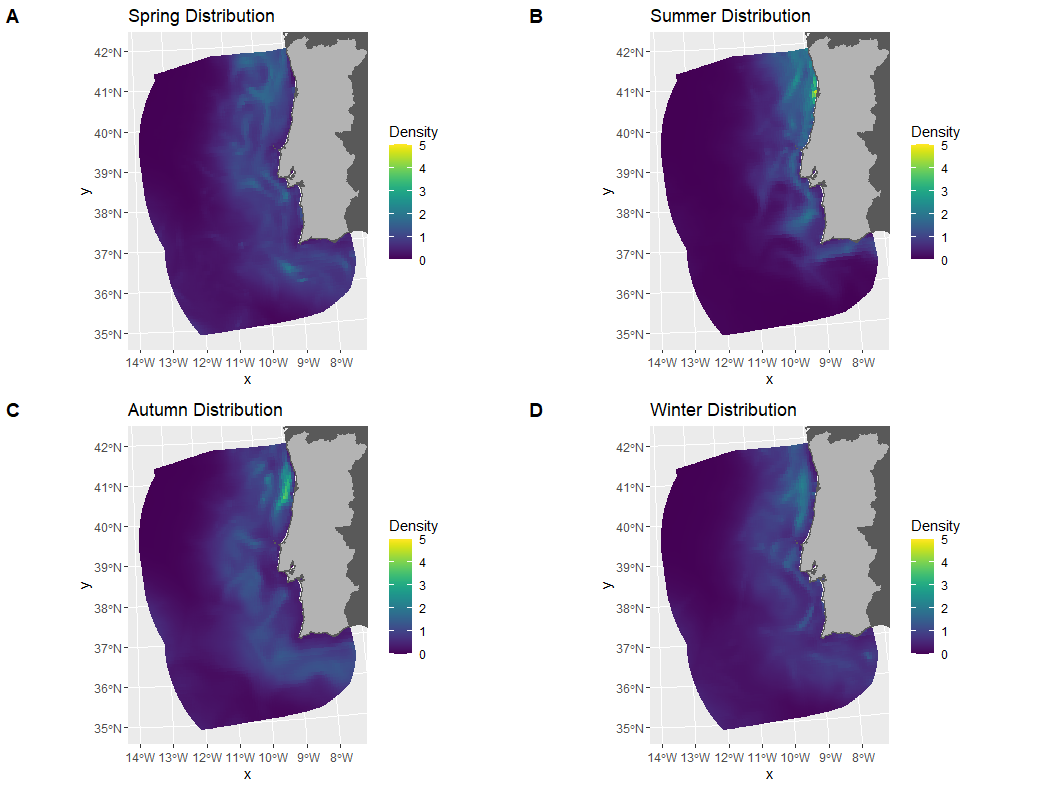

**FIGURE S19 -** Predicted common dolphin density distribution in mainland Portugal EEZ in 2018, according to the season.

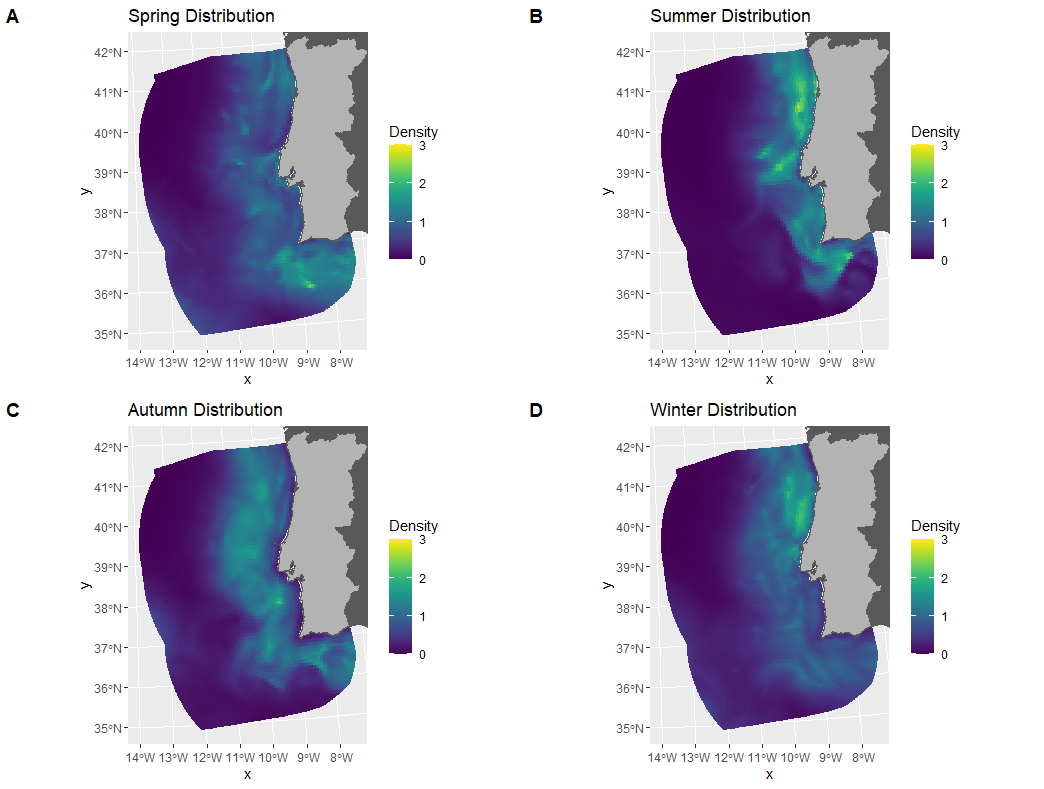

**FIGURE S20 -** Predicted common dolphin density distribution in mainland Portugal EEZ in 2019, according to the season.

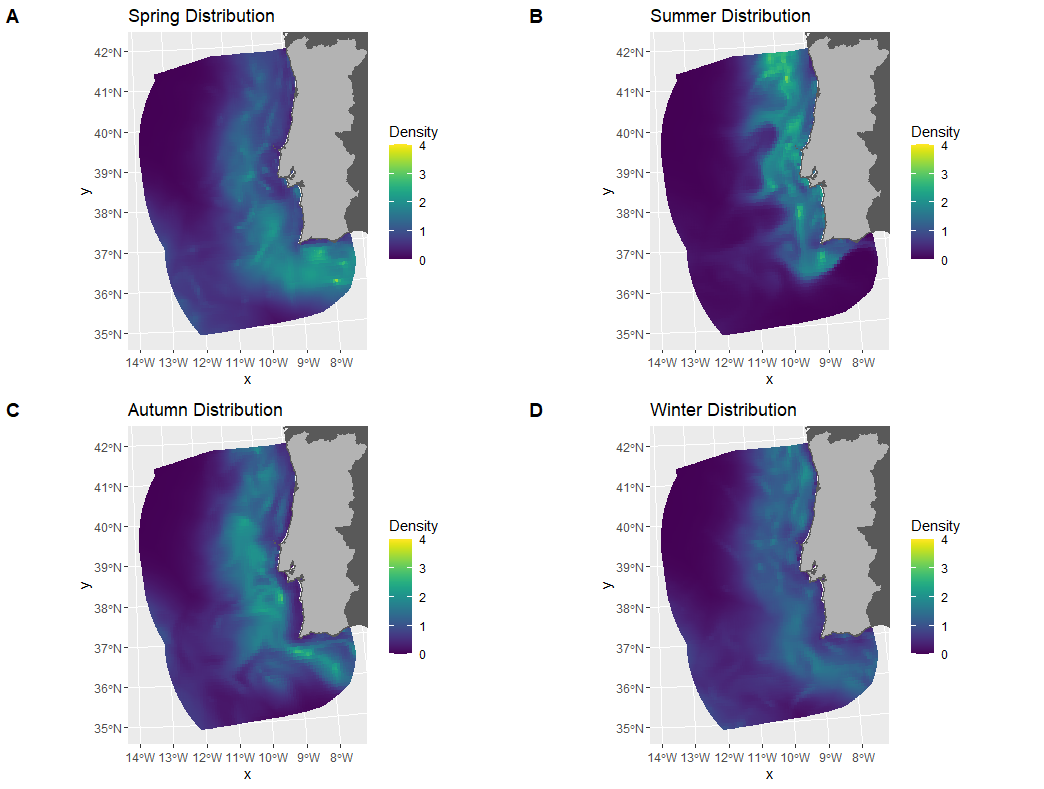

**FIGURE S21 -** Predicted common dolphin density distribution in mainland Portugal EEZ in 2020, according to the season.
